# Targeted Protein Relocalization via Protein Transport Coupling

**DOI:** 10.1101/2023.10.04.560943

**Authors:** Christine S. C. Ng, Aofei Liu, Bianxiao Cui, Steven M. Banik

## Abstract

Subcellular protein localization regulates protein function and can be corrupted in cancers^1^ and neurodegenerative diseases^2–4^. The localization of numerous proteins has been annotated^5–7^, and pharmacologically relevant approaches for precise rewiring of localization to address disease-driving phenotypes would be an attractive targeted therapeutic approach. Molecules which harness the trafficking of a shuttle protein to control the subcellular localization of a target protein could provide an avenue for targeted protein relocalization for interactome-rewiring therapeutics. To realize this concept, we deploy a quantitative approach to identify features which govern the ability to hijack protein trafficking, develop a collection of shuttle proteins and ligands, and demonstrate relocalization of proteins bearing endogenous localization signals. Using a custom imaging analysis pipeline, we show that endogenous localization signals can be overcome through molecular coupling of target proteins to shuttle proteins containing sufficiently strong native localization sequences expressed in the necessary abundance. We develop nuclear hormone receptors as viable shuttles which can be harnessed with Targeted Relocalization Activating Molecules (TRAMs) to redistribute disease-driving mutant proteins such as SMARCB1^Q318X^, TDP43 ^ΔNLS^, and FUS^R495X^. Small molecule-mediated relocalization of FUS^R495X^ to the nucleus from the cytoplasm reduced the number of cellular stress granules in a model of cellular stress. Using Cas9-mediated knock-in tagging, we demonstrate nuclear enrichment of both low abundance (FOXO3a) and high abundance (FKBP12) endogenous proteins via molecular coupling to nuclear hormone receptor trafficking. Finally, small molecule-mediated redistribution of NMNAT1 from nuclei to axons in primary neurons was able to slow axonal degeneration and pharmacologically mimic the WldS gain-of-function phenotype from mice resistant to certain types of neurodegeneration^8^. The concept of targeted protein relocalization could therefore nucleate approaches for treating disease through interactome rewiring.

## Introduction

Spatiotemporal control of subcellular protein localization creates a coordinated system to regulate cellular physiology. Aberrant trafficking and localization of proteins underlies numerous diseases, including cancers^9–11^, neurodegenerative diseases^3,12,13^, and genetic disorders^14,15^. Mutations which reroute tumor suppressor proteins from the nucleus to the cytoplasm are common mechanisms for promoting oncogenesis. Translocation of RNA-binding proteins such as TDP43 and FUS to the cytoplasm is a hallmark of amyotrophic lateral sclerosis (ALS)^2^ with numerous deleterious phenotypic consequences, including increased cytoplasmic aggregates^16–19^. Correcting these phenotypes through selective inhibition of protein trafficking would present several challenges given the conserved nature of nuclear export and import machinery. The global toxicity of nuclear import/export inhibition has restricted the application of molecules which target these pathways^20–22^. Yet, the clinical success of global nuclear export inhibitors such as Selinexor in certain contexts^23,24^ suggests that targeted approaches to control protein trafficking could find widespread use to reprogram protein localization.

The ability to exert targeted, pharmacological control of the subcellular location of an individual protein could provide avenues for addressing intractable diseases, as function-blocking molecules address only a small number of disease-relevant activities^25^. Gain-of-function molecules that directly drive the formation of neocomplexes between proteins lift several constraints that restrict the range of viable therapeutic targets^26–29^. In the context of controlling subcellular localization, correcting diseased phenotypes directly resulting from protein mislocalization or imparting beneficial function through protein relocation could expand the range of therapeutic options. A selection of tools for studying protein sequestration upon different stimuli with synthetic protein fusions to nuclear hormone receptor ligand binding domains ^30–32^, LOV2 domains^33–35^, or binding sites for localization sequence containing nanobodies^36^, have enabled fundamental studies of altered protein localization. Early work employing chemically-induced proximity between FRB and FKBP12 fused to arrayed localization sequences driven by rapamycin demonstrated the potential of small molecule control over localization programming^37–39^. This approach was extended to the rapid sequestration of cytoplasmic and endosomal proteins onto mitochondrial surfaces to attenuate activity^40^. Chemical control over localization has also been demonstrated with molecules which consist of protein-binding warheads linked to DNA intercalators^41^ and membrane-targeting lipids^42–44^.

Therapeutic modulation of target protein location via coupling to a cellular shuttle or anchor protein provides an alternative avenue for programmable molecular control. To advance the idea of localization control to address disease-driving proteins, an understanding of the cellular features which enable localization hijacking and expansion of the viable targets, molecules, and mechanistic principles of transport control are needed. A quantitative analysis of localization reprogramming ability would greatly facilitate extension of small molecule-mediated protein relocalization to therapeutic contexts. Here, we deploy a quantitative approach to demonstrate targeted protein relocalization via Targeted Relocalization Activating Molecules (TRAMs) which couple the trafficking of a target protein to the trafficking of a shuttle protein. We demonstrate the ability to employ nuclear hormone receptors to mediate nuclear import of targets including endogenous proteins and proteins underlying neurodegenerative diseases and cancers. Finally, we utilize targeted relocalization via a TRAM to generate a gain-of-function protective phenotype in a neuronal model of degeneration. Through coupling shuttles and targets using drug-like molecules, we advance approaches for therapeutic modulation of protein function based on the concept of controlled protein trafficking.

## Results

Nuclear localization and export sequences are recognized by members of the karyopherin family of shuttle proteins to regulate bidirectional transport of cargo between the nucleus and the cytoplasm^4^. We reasoned that to overcome the inherent localization of a protein via protein transport coupling would require 1) a sufficiently strong opposing localization sequence on the shuttling protein and 2) a requisite stoichiometry of shuttling protein. To examine these parameters, we chose nicotinamide nucleotide adenylyltransferase 1 (NMNAT1) as a model nuclear-localized target protein for relocalization. We hypothesized that since NMNAT1 possesses only a single predicted nuclear localization sequence (NLS) and no known direct DNA-binding domain, it might be susceptible to small molecule-mediated relocalization. To enable the study of proteins lacking known-small molecule binders, we synthesized the bifunctional molecule **1** (Fig.1b), which engages the *E*. *coli* dihydrofolate reductase domain (ecDHFR) and the FKBP12^F36V^ domain, both of which have found widespread use due to their tight binding affinities and orthogonality to most mammalian proteins. We reasoned that using non-covalent warheads would enable future extrapolation to the large number of available non-covalent and specific binders for proteins. We generated an mCherry fusion to ecDHFR and the nuclear export sequence from the HIV-1 rev protein^45^ to act as a shuttle protein. As transient transfection results in a range of confounding variables for examining the relative stoichiometries between two proteins and results in non-endogenous levels of transgene expression^46^, we stably incorporated mCherry-ecDHFR-NES and FKBP12^F36V^-GFP-NMNAT1 into HeLa cells (Fig. 1c). We isolated clonal export line A (**EL-A**) with a relative mCherry/GFP mean fluorescence intensity ratio of 1.0, which demonstrated a clear, dose-dependent translocation of NMNAT1 to the cytoplasm upon treatment with **1** (Fig. 1d), an effect that was abrogated at higher concentrations. Simultaneous treatment with the unlinked warheads **2** and **3** had no effect on protein localization (Extended Data Fig. 1a,b).

**Fig. 1:**
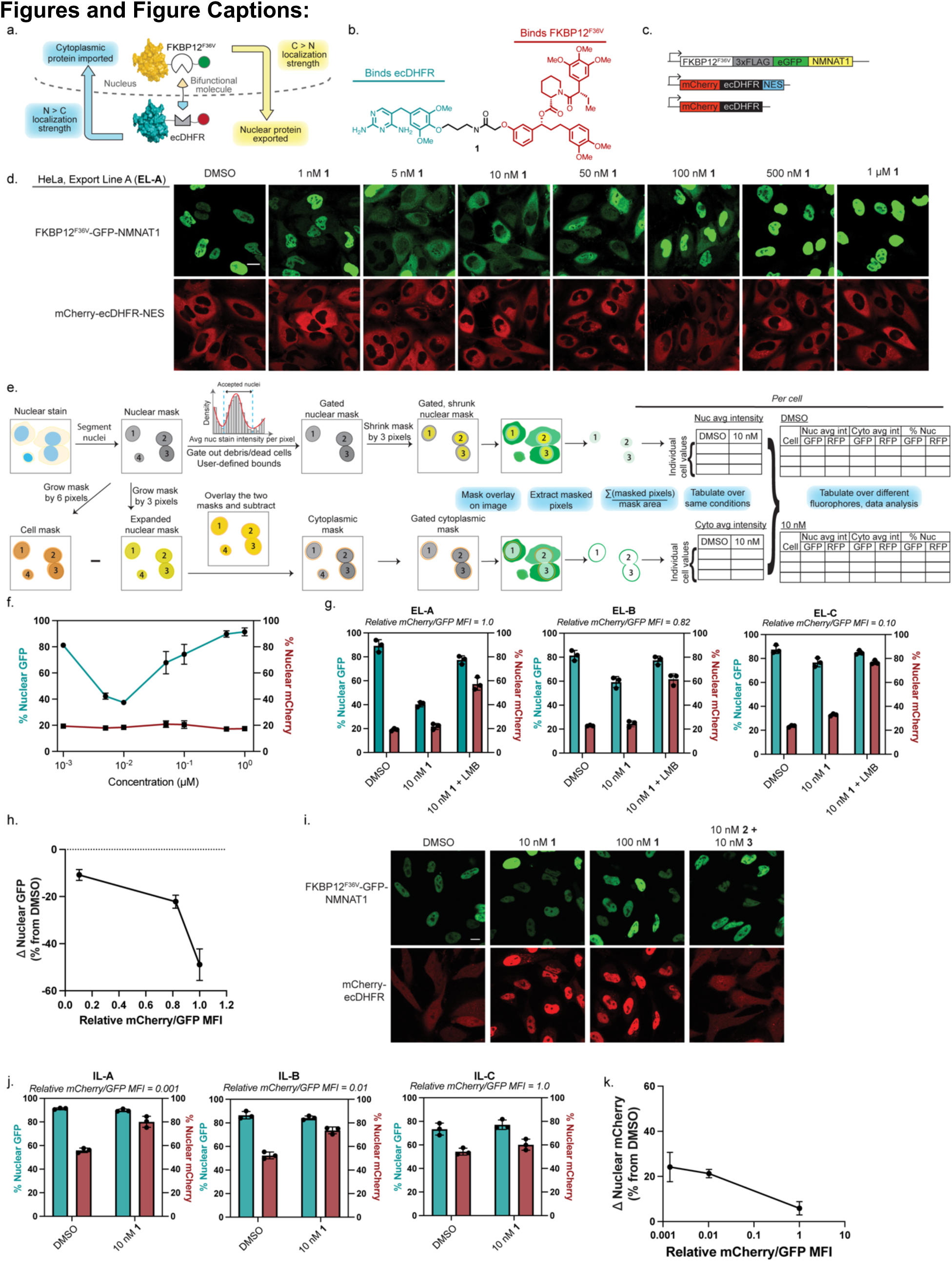
Development of a quantitative cell-by-cell analysis pipeline and application to targeted protein relocalization. **a**, Subcellular nucleocytoplasmic target localization control via protein transport coupling. **b**, Model bifunctional molecule which engages ecDHFR and FKBP12^F36V^ domains. **c**, Viral constructs for generating stable cell lines for studying localization control of NMNAT1. **d**, Representative live-cell images for NMNAT1 relocalization driven by **1** in Export Line A (EL-A) after 3 hour treatment. **e**, Schematic of computational pipeline for cell-by-cell imaging analysis of dual protein translocation. **f**, Molecule **1**-coupled nuclear export dose response derived using analysis pipeline in e after 3 hour treatment. **g**, Ability of three NMNAT1 export lines exhibiting different relative export shuttle (mCherry) to NMNAT1 (GFP) ratios to translocate NMNAT1 from the nucleus promoted by **1**, or inhibited by leptomycin B (LMB) treatment after 3 hour treatment. **h**, Impact of relative stoichiometry on nuclear export of NMNAT1. **i**, Representative images of nuclear import of a diffuse protein driven by NMNAT1 after 3 hour treatment with **1**. **j**, Ability of three representative import cell lines (IL) with different relative target (mCherry) and shuttle (NMNAT1) ratios to redistribute a diffuse protein target after 3 hour treatment with **1**. **k**, Impact of relative stoichiometry on nuclear import of mCherry. Images in **d, i** are representative of three biological replicates. Data are compiled from three independent experiments. Scale bars are 20 µm.

The quantification of nucleocytoplasmic transport when studying two proteins simultaneously has traditionally been done by a colocalization metric across a set of representative images. We reasoned that an open-source computational imaging analysis pipeline that provides single cell-level analysis of two different proteins simultaneously in both the nucleus and cytoplasm would be the most accurate approach for gauging protein transport and comparing different target proteins and shuttle proteins (Fig. 1e). We built this custom analysis pipeline by first segmenting cells using a nuclear stain reference, then masking cells to capture both a nuclear area and cytoplasmic area using a flexible pixel distance definition. We applied a gating filter on nuclear intensities to remove aberrantly high nuclear-stained objects such as debris and dead cells. Finally, we summed and averaged the intensities per cell for the nuclear masked area and cytoplasmic masked area to give a pixel intensity per area measurement akin to the relative concentration of fluorophore in the nucleus and cytoplasm for each cell. We verified this approach would be applicable to numerous cell types and densities (Extended Data Fig. 2a-c.).

We used our quantitative analysis pipeline to examine dose-, shuttle protein- and stoichiometry-dependent relocalization. We observed a concentration-dependent hook effect with **1** on the transport of NMNAT1 (Fig. 1f). Quantification of transport also revealed that while NMNAT1 was readily extracted to the cytoplasm with **1**, NMNAT1 did not possess a sufficiently strong NLS to partially redistribute mCherry-ecDHFR-NES to the nucleus. To examine the impact of relative stoichiometry between NMNAT1 and ecDHFR-NES, we isolated two additional clonal export cell lines (**EL-B**,**C**) with varying mCherry:GFP ratios (Extended Data Fig. 1c). When treated with **1**, cell lines with higher mCherry:GFP ratios exhibited a high degree of NMNAT1 redistribution to the cytoplasm (Fig. 1g). Cell lines with intermediate ratios resulted in a diffusive state where NMNAT1 was found both in the cytoplasm and nucleus. Cells with the lowest relative expression levels of mCherry to GFP (**EL-C)** exhibited only 10% NMNAT1 relocalization upon treatment with **1** (Fig. 1h). When cells were cotreated with leptomycin B (LMB), an inhibitor of nuclear export by exportin 1, along with **1**, no relocalization of NMNAT1 was observed and an enrichment in nuclear mCherry resulted (Extended Data Fig. 1d-f). Similar to **EL-A**, cotreatment with **2** and **3** did not cause protein relocalization in **EL-B**,**C** (Extended Data Fig. 1d-f). No degradation was observed as a function of **1** in four clonal cell lines (Extended Data Fig. 1g,h). These results support active NES-driven redistribution of NMNAT1 through small molecule-mediated coupling, and the concept of relative localization strength hierarchy where a single NES from a shuttle protein can overcome the native NLS of a target protein.

Several proteins inherently localize simultaneously to both the nucleus and cytoplasm. To examine if diffuse proteins can be sequestered in the nucleus by non-DNA binding proteins, we generated cell lines stably expressing mCherry-ecDHFR without an NES as well as FKBP12^F36V^-GFP-NMNAT1. We isolated clonal import cell lines A-C (**IL-A–C**) with relative mCherry:GFP ratios spanning three orders of magnitude (Extended Data Fig. 2a) and analyzed their ability to sequester mCherry upon treatment with **1**. The initial percentage of nuclear mCherry was ∼50% in all cell lines, consistent with diffuse localization. After treatment with **1**, lines **IL-A** and **IL-B** with relative mCherry:GFP ratios of 0.03 and 0.1, respectively, were both capable of significant sequestration of mCherry (80% nuclear) (Fig. 1i,j, Extended Data Fig. 2b-d). Cotreatment with **2** and **3** had no effect on mCherry localization (Extended Data Fig. 2b,c). **IL-C**, with a relative mCherry:GFP ratio of 1.0 did not exhibit a large enrichment of nuclear mCherry upon treatment with **1** (Fig. 1j,k, Extended Data Fig. 2e). No diffusion of FKBP12^F36V^-GFP-NMNAT1 was observed in cells because of coupling to mCherry-ecDHFR (Extended Data Fig. 2f). Treatment with **1** did not lead to degradation of either mCherry or NMNAT1 in four clonal cell lines (Extended Data Fig. 2g,h). Together, these data demonstrate that sufficient levels of anchor or shuttle must be present to enforce full sequestration of a target protein.

In principle, the trafficking of any protein might serve to relocalize another provided sufficient localization strength, expression levels, and effective molecular coupling. We reasoned that ideal nuclear shuttle proteins which transport cargo from the cytoplasm to the nucleus might be proteins which are capable of traversing both compartments, but upon ligand recognition, localize strongly to the nucleus. Nuclear hormone receptors such as the estrogen receptor (ERα) and glucocorticoid receptor (GR) exhibit ligand-dependent translocation from the cytoplasm to the nucleus to exert downstream transcriptional activation (Fig. 2a). Several small molecule inhibitors have been developed to inhibit the transcriptional functions of nuclear hormone receptors which might be repurposed to hijack the nuclear trafficking capabilities of these proteins. To demonstrate the potential of these receptors as nuclear shuttles, we synthesized **4** and **5** (Fig. 2b), which utilize a raloxifene-based warhead known to engage ERα but not lead to its downregulation.

**Fig. 2:**
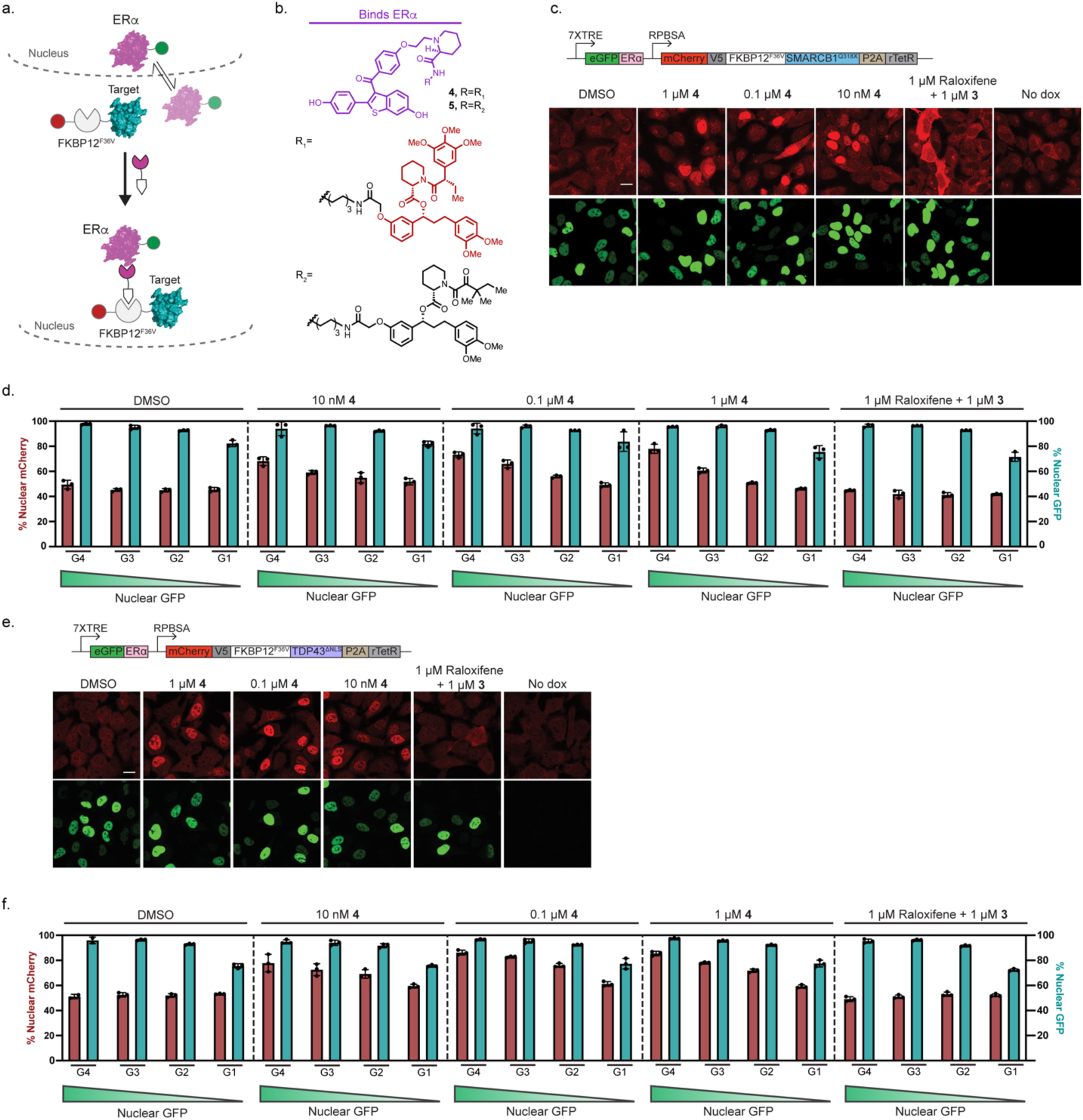
Nuclear hormone receptors can act as shuttles for cytoplasmic and diffuse proteins. **a**, Schematic for ERα dependent nuclear import of target proteins. **b**, Bifunctional molecules which engage ERα and FKBP12^F36V^ or FKBP12. **c**, Genetic constructs for generating stable lines and representative images of HeLa cells stably expressing the SMARCB1^Q318X^ mutant and ERα under an inducible promoter showing import of SMARCB1^Q318X^ after a 3 hour treatment with **4**. **d**, Quantitative group analysis of cell distribution expressing mCherry-SMARCB1^Q318X^ and GFP-ERα after a 3 hour treatment with **4**. **e**, Representative images of HeLa cells stably expressing the TDP43 ^ΔNLS^ mutant and ERα under an inducible promoter, showing import of SMARCB1^Q318X^ after a 3 hour treatment with **4**. TDP43 ^ΔNLS^ = TDP43^(K82A/R83A/K84A)^. **f**, Quantitative group analysis of cell distribution expressing mCherry-TDP43 ^ΔNLS^ and GFP-ERα, For **d** and **f**, Cells were divided into 4 groups based on their average nuclear GFP intensity G4: 4500 > x ≥ 4500, G3: 3000 > x ≥ 1500, G2: 1500 > x ≥ 500, G1: 500 > x ≥ 50.

Mislocalized transcriptional regulators and RNA-binding proteins are hallmarks of several cancers and neurodegenerative diseases. Targeted approaches which return these proteins to the nucleus might help ameliorate disease-driving phenotypes. We first examined the ability to nuclear enrich SMARCB1^Q318X^, a mutant C-terminal truncation of SMARCB1 found in atypical teratoid/rhabdoid tumors which has been reported to exhibit cytoplasmic localization^47^. We engineered HeLa cells to constitutively express SMARCB1^Q318X^ along with ERα on a doxycycline inducible promoter (Fig. 2c). SMARCB1^Q318X^ exhibited diffuse localization, which upon addition of doxycycline and subsequent treatment with **4** could be enriched in the nucleus (Fig. 2c). To assess the impact of ERα-GFP stoichiometry on nuclear enrichment, we analyzed cells grouped based on relative ERα-GFP expression levels (Fig. 2d). In the grouping with the highest ERα expression, nuclear relocalization was observed at all concentrations of **4**, with the most substantial redistribution occurring at 1 µM (80% nuclear SMARCB1^Q318X^). Within the lowest expression ERα-GFP grouping (G1), no relocalization was observed. Across all concentrations of **4**, it was evident that high expression of the nuclear shuttle protein drove the ability to redistribute SMARCB1^Q318X^. Finally, even at low levels of ERα-GFP, no nuclear exclusion was observed as a function of treatment with **4**, suggesting SMARCB1^Q318X^ is incapable of overcoming the trafficking activity of ERα in a reciprocal fashion.

Mislocalized TDP43 mutants which seed aggregates in the cytoplasm are a conserved biomarker for ALS.^48,49^ We applied an analogous approach to that of SMARCB1^Q318X^ to examine TDP43 ^ΔNLS^ relocalization back to the nucleus using ERα. In the absence of small molecule treatment, we observed diffuse TDP43 ^ΔNLS^ (Fig. 2e). Upon treatment with **4**, we observed significant nuclear enrichment across concentrations spanning 10 nM–1 µM in the top ERα-GFP expressing groups (Fig. 2f). Together, these results demonstrate that sufficient expression of the shuttle protein is essential for achieving targeted protein redistribution and that truncated mutant pathogenic proteins are susceptible to small molecule-induced relocalization via transport coupling.

Mutations in the Fused in Sarcoma (FUS) protein which alter its nuclear localization have been observed in ALS patients.^50,51^ The FUS^R495X^ mutation is associated with an aggressive disease phenotype^52,53^, and forms cytoplasmic granules in neurons which have been linked to its pathogenic function.^54,55^ To examine the ability for transport-coupled relocalization to redistribute FUS^R495X^ from the cytoplasm to the nucleus, we generated HeLa cells which stably express mCherry-FKBP12^F36V^-FUS^R495X^ and ERα-GFP under a doxycycline inducible promoter (Fig. 3a). Upon induction of ERα expression and treatment with **4**, we observed significant redistribution of FUS^R495X^ from the cytoplasm to the nucleus, which did not occur when cells were co-treated with raloxifene and **3**, or in the absence of ERα (Fig. 3a). Analysis of cell groups with different ERα expression levels showed both dose-dependence and significant relocalization (up to 90%) across ERα expression levels (Fig. 3b). We hypothesized that FUS^R495X^-positive cytoplasmic granules might be dissolved if targeted protein relocalization could redistribute FUS^R495X^ to the nucleus. To assess the ability to extract proteins from granules, we examined FUS^R495X^-expressing HeLa cells induced with doxycycline and treated with sodium arsenite to instigate stress granule formation (Fig. 3c). Upon formation of granules, cells were treated with **4** and continuously imaged. We observed relocalization of FUS^R495X^ from both granules and the cytoplasm to the nucleus over the course of two hours after exposure to **4** (Fig. 3d). These observations are consistent with small molecule-mediated active extraction of FUS^R495X^ from cytoplasmic granules to the nucleus. To quantify the effect of FUS^R495X^ redistribution on the number of stress granules in cells, we performed fixed-cell immunofluorescence imaging to examine both mCherry-FKBP12^F36V^-FUS^R495X^ and G3BP1, a cellular marker of stress granules (Fig. 3e). We observed a significant reduction in both FUS^R495X^-positive and G3BP1-positive stress granules in cells treated with **4** (Fig. 3f). We also observed a reduction in both FUS^R495X^-postive and G3BP1-positive stress granules in cells cotreated with raloxifene and **3**, likely through small-molecule induced stabilization of the FKBP12^F36V^ domain of FUS^R495X^ by **3**. However, treatment with **4** led to more substantial reductions (Fig. 3g,h), and demonstrates that relocalization of FUS^R495X^ from stress granules to the nucleus can result in dissolution of FUS^R495X^-positive and G3BP1-positive granules.

**Fig. 3:**
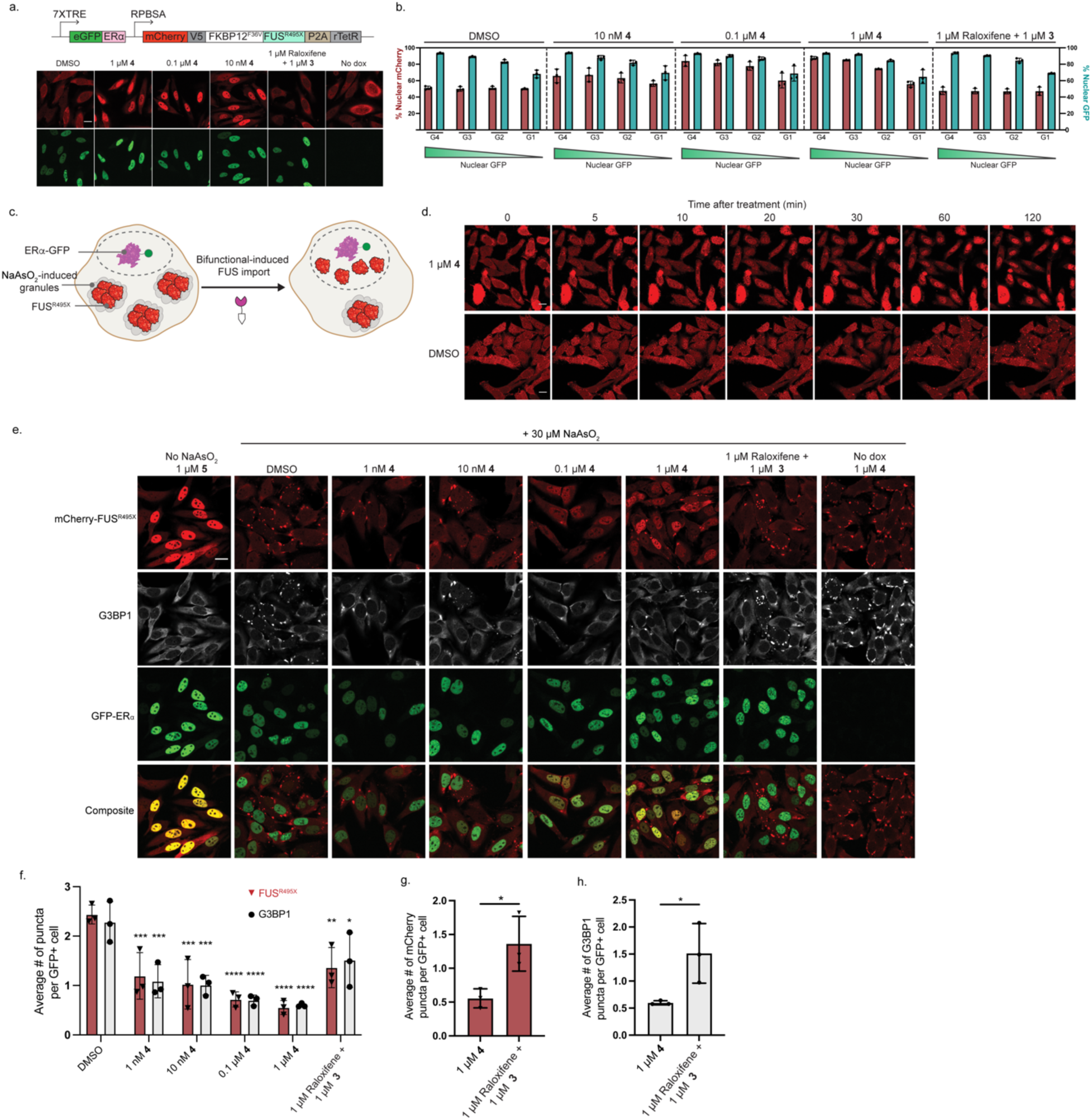
Small molecule-mediated relocalization can extract proteins from stress granules. **a**, Representative images of HeLa cells stably expressing a FUS^R495X^ mutant and ERα under an inducible promoter. **b**, Quantitative group analysis of cell distribution expressing mCherry-FUS^R495X^ and GFP-ERα. Cells were divided into 4 groups based on their average nuclear GFP intensity G4: 1500 > x ≥ 800, G3: 800 > x ≥ 500, G2: 500 > x ≥ 200, G1: 200 > x ≥ 50. **c**, Schematic for ERα-induced relocalization of FUSR495X from stress granules formed by exposure to NaAsO_2_. **d**, Timelapse live-cell imaging snapshots of FUS^R495X^ extraction from granules after treatment with **4**. **e**, Representative fixed-cell immunofluorescent images of stress granule marker G3BP1 and mCherry-FUS^R495X^ after treatment with **4** or control compounds. **f**, Quantification of stress granules in cells after treatment with varying concentrations of **4** or control compounds. **g**, Comparison of mCherry-FUS^R495X^ positive puncta in cells treated with bifunctional molecule **4** or unlinked control molecules. **h**, Comparison of G3BP1-positive puncta in cells treated with bifunctional molecule **4** or unlinked control molecules. Scale bars are 20 µM. Images in **a** and **e** are representative of three independent experiments. Images in **d** are representative of two independent experiments. Three fields of view totaling to a minimum of 40 cells per condition per repeat were used to generate mean values for **f**, **g**, **h**. P values in **f** were determined by two-way ANOVA comparing each dataset to its relative DMSO control. P values in **g**, **h** were determined by unpaired two-tailed t-tests. Data in **f**, **g**, **h** are shown as the mean of three independent experiments ± SD values. P values: (*) indicates P ≤ 0.05 (**) indicates P ≤ 0.01, (***) indicates P ≤ 0.001, P ≤ 0.0001.

The examination of targeted relocalization of endogenous proteins as a therapeutic approach is limited by a lack of small molecule ligands for any desired target protein. To bypass this challenge, we utilized a modified Cas9-based protocol for microhomology-mediated end joining to install binding domains and fluorescent proteins on endogenous protein targets (PITCh)^56^ (Fig. 4a). We redesigned the PITCh guide RNA plasmid to have two orthogonal U6 promoters and stem loops to facilitate rapid cloning for knock-in onto different targets. To examine the ability of ERα and GR to relocalize endogenous proteins, we first inserted FKBP12^F36V^-GFP onto FOXO3A in HEK293T cells (Fig. 4b). FOXO3A is a tumor suppressor transcription factor which is sequestered in the cytoplasm in numerous cancers^57,58^. We expressed ERα in our HEK293T FOXO3a-FKBP12^F36V^-GFP cells, and upon treatment with **4**, observed redistribution of FOXO3A to the nucleus in ERα-expressing cells (Fig. 4c). Only in the cell groups expressing the highest levels of ERα was significant redistribution observed, demonstrating the potential of molecular coupling to ERα to overcome the endogenous localization programming of cytoplasmic FOXO3A (Fig. 4d).

**Fig. 4:**
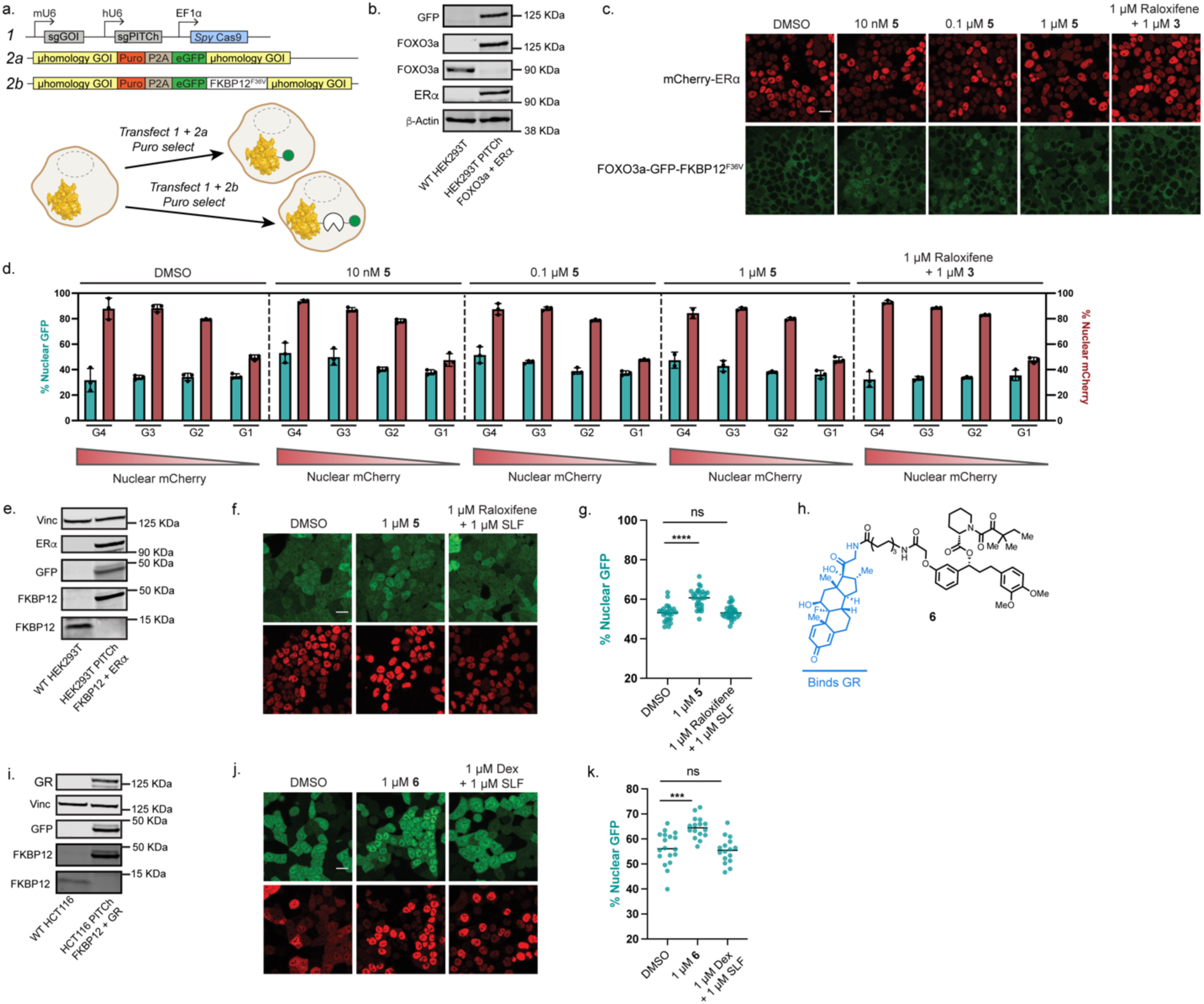
Endogenously-tagged proteins are susceptible to relocalization using nuclear hormone shuttles. **a**, Schematic for modified PITCh knock-in into target proteins. **b**, Demonstration of GFP-FKBP12^F36V^ tagging of endogenous FOXO3A in HEK293T cells stably expressing ERα. **c**, Representative live-cell images of FOXO3A knock-in HEK293T cells containing ERα and treated with **4** or small molecule controls. **d**, Group quantitative analysis of FOXO3A knock-in cells treated with varying concentrations of **4** for 3 hours. Cells were divided into 4 groups based on their average nuclear mCherry intensity G4: 4900 > x ≥ 2000, G3: 2000 > x ≥ 800, G2: 800 > x ≥ 200, G1: 200 > x ≥ 50. **e**, Demonstration of GFP tagging of endogenous FKBP12 in HEK293T cells stably expressing ERα. **f**, Representative live-cell images of FKBP12 knock-in HEK293T cells containing ERα and treated with **5** or small molecule controls for 3 hours. **g**, Quantitative analysis of nuclear FKBP12 in the top 5% ER-expressing HEK293T cells treated with **5** or small molecule controls for 3 hours. Number of cells: DMSO: 26, 1 μM **5**: 28, 1 μM Raloxifene + 1 μM SLF: 30. **h**, Bifunctional molecule which engages GR and FKBP12. **i**, Demonstration of GFP tagging of endogenous FKBP12 in HCT116 cells stably expressing GR. **j**, Representative live-cell images of FKBP12 knock-in HCT116 cells containing GR and treated with **6** or small molecule controls for 3 hours. **k**, Quantitative analysis of nuclear FKBP12 in the top 5% GR-expressing HCT116 cells. Number of cells: DMSO: 18, 1 μM **6**: 16, 1 μM Dexamethasone + 1 μM SLF: 16. Scale bars are 20 µm. Immunoblots are representative of two independent experiments. Images are representative of three biological replicates. Data are compiled from three independent experiments. P values in **g**, **j** were determined by unpaired two-tailed t-tests of the respective datasets to DMSO. P values: (*) indicates P ≤ 0.05 (**) indicates P ≤ 0.01, (***) indicates P ≤ 0.001, P ≤ 0.0001.

To explore the ability to shift the distribution of abundant, non-localized proteins, we also tagged endogenous FKBP12 with GFP in HEK293T cells stably expressing mCherry-ERα (Fig. 4e). We observed nuclear concentration of FKBP12 in cells treated with **5** (Fig. 4f, Extended Data Fig. 9a), but found no significant relocalization using cell group analysis (Extended Data Fig. 9b,c). We reasoned that high expression of FKBP12 might necessitate analysis of only the highest shuttle receptor-expressing cells. When we analyzed the top 5% of ERα-expressing cells, we found significant nuclear concentration of endogenous FKBP12 as a function of treatment with **5** (Fig. 4g, Extended Data Fig. 9d). To assess the ability to harness the glucocorticoid receptor (GR) as a nuclear shuttle, we synthesized **6** which utilizes dexamethasone as a GR-binding warhead (Fig. 4h). We also endogenously tagged FKBP12 in HCT116 cells, and stably incorporated GR as an mCherry fusion (Fig. 4i). Similar to HEK293T cells, we observed nuclear concentration of FKBP12 in HCT116 cells which was not significant by cell group analysis (Fig. 4j, Extended data Fig. 10a-c). However, significant nuclear concentration of endogenous FKBP12 was observed in the top 5% of cells with the highest expression of GR (Fig. 4k, Extended Data 10d). Cotreatment with SLF and dexamethasone did not result in nuclear concentration of FKBP12 (Extended Data Fig. 10b,c). We also observed ligand-dependent concentration of GR in the nucleus (Extended Data Fig. 10c), consistent with a ligand-gated TRAM for protein relocalization. We also generated the HCT116 endogenously-tagged FKBP12 line expressing mCherry-ERα, and observed no significant relocalization of FKBP12 at any expression level (Extended Data Fig. 11). We compared the relative expression levels of mCherry-GR and mCherry-ERα, and observed higher expression of mCherry-GR in our stably expressing HCT116 cells (Extended Data Fig. 10e). As FKBP12 is expressed at higher levels than FOXO3A by transcriptomic analysis, these data support the importance of relative stoichiometries between target and shuttle proteins when assessing targeted protein relocalization (Extended Data Fig. 9e, 10e). Further, the ability to analyze cell populations with specific shuttle-protein expression levels demonstrates the utility of a quantitative analysis pipeline.

Targeted protein relocalization via coupling to a shuttle protein could enable pharmacological approaches which mimic beneficial gain-of-function mutations. WldS, a mutant protein consisting of mouse NMNAT1 fused to Ube4B has been shown to protective against Wallerian neurodegeneration^59–61^. Mice bearing the WldS mutation have shown increased resistance to neuropathies and ALS. The ability for small amounts of WldS to traffic to the axon is crucial for its protective function, where it serves to maintain the levels of NAD^+^ upon loss of axonal NMNAT2 during early stages of axonal injury^62^. We reasoned that small molecule-mediated transport of NMNAT1 from the nucleus down the axon might serve a similar function to WldS and provide a proof-of-concept example for protein relocalization to drive a gain-of-function beneficial phenotype. We harvested DRG neurons from rat embryos and transduced them with AAVs expressing mouse NMNAT1 (mNMNAT1) linked to FKBP12^F36V^ and a truncated sequence of GAP-43, which localizes in axons, fused to ecDHFR (Fig. 5b). When we treated these neurons with **1**, clear redistribution of NMNAT1 down axons was observed, which did not occur when neurons were cotreated with **2** and **3** (Fig. 5c). To assess the axonal protective function of NMNAT1 redistribution, after treatment with **1**, we performed an axotomy to remove the cell bodies. We observed persistent mNMNAT1 expression in axons post axotomy only in neurons treated with **1** (Fig. 5d). Distinct blebbing and fragmentation of neurites was observed in DMSO-treated neurons 12-24 hours post axotomy, with near complete neurite death observed after 48 hours (Fig. 5e). However, in neurons treated with **1**, both axon and axon termini health were maintained for at least 48 hours post-axotomy and attenuated axon death observed only after 96 hours. These observations are consistent with gain-of-function protective activity of NMNAT1 driven by targeted, small molecule-mediated relocalization from nuclei to axons.

**Fig. 5:**
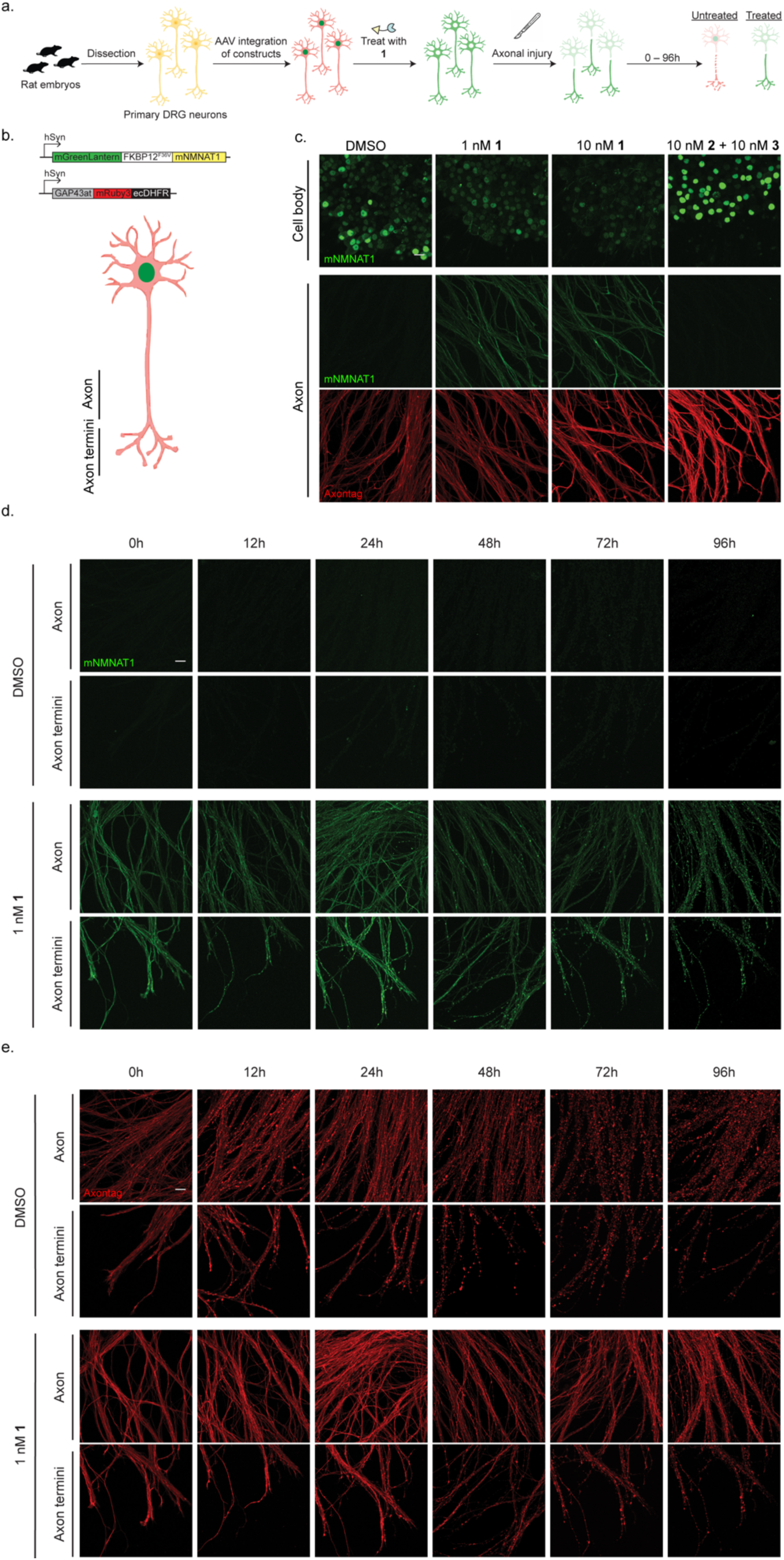
Targeted relocalization of NMNAT1 can protect against axon injury. **a**, Schematic for DRG neuron harvesting, transduction of model proteins via AAV, small molecule treatment and axotomy. **b**, Constructs delivered by AAV to study the gain-of-function potential for mNMNAT1 relocalization in neurons, and definition of labels for neuronal imaging. GAP43at is the 20 amino acid dipalmitoylation domain (MLCCMRRTKQV) found at the N terminus of growth-associated protein-43. **c**, Representative images of mNMNAT1 in primary DRG neurons when treated with **1** or control molecules. **d**, Representative images of mNMNAT1 in axons after exposure to **1** for 24 hours followed by axotomy to induce degeneration. **e**, Representative images of axonal-tagged protein in axons and axon termini after treatment with **1** for 24 hours and axotomy. mRuby3 signal was used to monitor degeneration across treated neurons. Axontag = GAP43at-mRuby3-ecDHFR. Images are representative of neurons harvested from two different embryos. Scale bars are 20 µm.

## Discussion

The onset of strategies which utilize endogenous cellular machinery to manipulate or control therapeutic targets has instigated exploration of new ways to rewire cellular biology. Subcellular protein localization often serves as a control mechanism for protein activity, and alterations in a protein’s localization can form the basis of numerous diseases. Controlling a protein’s location through hijacking the transport mechanisms of another protein could serve as a viable therapeutic approach for precision manipulation of protein function. In the present study, we develop a quantitative single-cell analysis approach for examining the potential for targeted relocalization-activating molecule (TRAM)-mediated coupling of a target protein with a shuttle protein. We discover that the relative expression levels of the two protein partners is a predominant parameter in driving the ability to relocalize a target protein. We also demonstrate that the endogenous localization of a target protein can be overturned by a sufficiently strong opposing localization signal and levels of the shuttle protein. Pioneering work using chemical methods to influence nucleocytoplasmic transport have relied on pitting two synthetic localization signals against each other^37^. Extending these efforts to native systems, as we do in the present study, represents the next frontier in controlling mammalian protein localization pharmacologically. How to quantify relative localization strength between proteins remains an open question^63–65^, with context dependencies that are likely intrinsic to each individual protein. Quantitative analysis at the single-cell level is essential for assessing the ability for small molecules to coopt other mammalian machinery.

Tethering two proteins together to alter the trafficking of one partner necessitates suitable shuttle proteins with ligands. Nuclear hormone receptors are common targets for small molecule therapeutics and exhibit nuclear translocation or localization. We have developed molecules which employ binders of nuclear hormone receptors and can program them to drive enrichment of a protein target to the nucleus. Utilizing discrete shuttles that are differentially expressed in disease contexts such as cancer or uniquely expressed in certain cell types presents the future prospect of targeted relocalization-based therapies which can generate beneficial phenotypic consequences^66–69^.

The mislocalization of transcriptional regulators or RNA-binding proteins has been observed in cancers and neurodegenerative diseases. Mutations which truncate or impair nuclear localization sequences are particularly pathogenic, with no clear mechanism for therapeutic intervention beyond genetic manipulation. We demonstrate the ability of small molecule-mediated targeted relocalization utilizing nuclear hormone receptors to return these proteins to the nucleus from the cytoplasm or stress granules. Redistribution of mislocalized proteins from stress granules back to the cytoplasm might serve to both restore native function and protect against pathogenic function. Our data suggests that the redistribution of mislocalized proteins which form the core of stress granules back to nuclei might dissipate these granules.

Targeted redistribution of endogenous proteins driven by small molecules has been challenging to rigorously demonstrate due to both a lack of small molecule ligands and methods for imaging and analyzing protein localization quantitatively with high sensitivity. We show, via endogenous knock-in of binding domains or fluorescent proteins, the ability to redistribute endogenous proteins to the nucleus from the cytoplasm using nuclear hormone receptor shuttles. We demonstrate that the native NES of FOXO3A can be overcome by coupling to ERα. Using SLF, we also demonstrate concentration of FKBP12 in the nucleus. FKBP12 is expressed at high levels in the cell lines utilized in the present study, and only in the cells expressing the highest levels of shuttle protein was nuclear concentration observed, highlighting the importance of matching shuttles and targets to achieve the desired degree of protein movement.

An attractive aspect of targeted protein relocalization is the ability to impart protein function in a new subcellular compartment. The WldS mutation has been extensively studied for its ability to confer protection against neurodegenerative conditions. A consensus protective function of WldS is maintaining the balance of NAD^+^ in axons during injury. Small molecule approaches to target this axis have attempted to inhibit the widely expressed NADase enzyme SARM1^70,71^. Targeted relocalization of NMNAT1 might serve to compensate for the loss of NMNAT2 during early neurodegeneration and could present an alternative approach amenable to specific targeting using neuron-specific transport pathways. We demonstrate the ability to utilize such pathways for moving nuclear proteins down axons in a targeted fashion.

While we employ genetic-fusion constructs to examine the relocalization of protein targets, these studies motivate further in-depth ligand development campaigns which disclose non-inhibitory small molecules. The degree to which any individual target needs to be redistributed to achieve a phenotypic effect is likely to be context dependent. Thus, we anticipate interactome rewiring for manipulating protein localization has the potential to contribute to advances in understanding both fundamental biology and developing next-generation therapeutics.

## Acknowledgements

We thank Dr. Melissa Gray for experimental assistance. This work was supported by an A*STAR fellowship to C.S.C.N.

## Author Contributions

C. S. C. N. and S.M.B. conceived of the project. C. S. C. N. conducted all experiments and analyzed all data. A. L. and B.C. provided essential expertise on neuron harvesting and biology. C.S.C.N and S.M.B. wrote the manuscript. S.M.B. provided supervision.

## Methods

### Cell lines

Adherent cells were cultured in 10 or 15 cm plates at 37 °C in a 5% CO_2_ humidified atmosphere. HeLa, HCT116, and HEK293T cells were obtained from ATCC and were cultured in DMEM supplemented with 10% heat-inactivated fetal bovine serum (HI-FBS) and 1% penicillin/streptomycin.

### Stable Cell Line Generation with Sleeping Beauty Transposase

Fusion protein constructs were cloned into a pSBtet vector flanked by Sleeping Beauty transposase recognition sites which also encoded constitutive expression of the puromycin resistance gene. The day before transfection, 300,000 cells were seeded into a 6-well plate. The next day, 1.9 µg of transgene DNA was mixed with 0.1 µg of Sleeping Beauty Transposase encoding plasmid in 200 µL of JetPrime buffer, then 4 µL of JetPrime reagent was added. The mixture was gently mixed and left to incubate for 10 minutes at room temperature, after which it was added dropwise to the well. The next morning, media was replaced with fresh cell culture medium. Three days post transfection, antibiotic selection was initiated with 1 µg/mL puromycin until all control cells had died.

### Retrovirus Generation

One day before transfection, 1.5 million HEK293T cells were seeded into 6 cm dishes. The following day, 130 µL complete DMEM media was added into an Eppendorf tube. 1 µg of transgene cloned into a pMXs vector was added along with 900 ng of the retrovirus pol/gag and 150 ng of VSVg DNA, and gently mixed. PEI (1 mg/mL) was brought to room temperature and gently mixed before adding 6 µL to the DNA mixture. The mixture was gently mixed and incubated for 20 minutes at room temperature. Media in the 6 cm dishes was replaced with fresh complete DMEM. After 20 minutes incubation, the transfection mixture was added dropwise to the 6 cm plate, then the plate was returned to the incubator. The next morning, media was removed and replaced with 5 mL DMEM media containing 30% heat-inactivated fetal serum and 1% Pen/Strep and the plate was returned to the incubator. After 36 hours, the media from the 6 cm plate was collected into 15 mL falcon tube and centrifuged at 1000 RPM for 5 minutes. The supernatant was filtered through a 0.45 µM filter into 300 µL aliquots. Aliquots were stored at -80 °C.

### Generating Stable Cell Lines with Retrovirus

On day of infection, cells were seeded at a density of 2 million cells per well in a 6 well plate, and media in the well was added up to 3 mL total volume. 3 µL of polybrene transfection reagent (EMD Millipore Corp. #TR-1003-G) was added to each well. Viral aliquots were thawed at 37 °C, then gently mixed and added to the well. The plate was the centrifuged at 2200 RPM for 45 min at 37 °C, then returned to the incubator. The next morning, the cells were lifted and transferred into a 10 cm plate. When cells reached 70-80% confluency, antibiotic selection was started with 10 µg/mL blasticidin until control cells died.

### Lentivirus Generation

1.2 million viral HEK293T cells were seeded per well in a 6 well plate. The next day, 6 µL Mirus-LT1 transfection reagent was mixed with 250 µL OptiMEM and left to incubate at room temperature for 15 minutes. 1.5 µg transgene cloned into the pU6 transfer vector was mixed with 0.1 µg gag/pol, 0.1 µg REV, 0.1 µg TAT and 0.2 µg VSVg packaging plasmids. The OptiMEM and Mirus-LT1 mixture was then added to the DNA mixture, mixed and left to incubate at room temperature for 15 minutes, then added dropwise to well. Plate was gently swirled to mix, then returned to the incubator. The next morning, media in the well was replaced. 36 hours later, the supernatant was retrieved into 15 mL falcon tubes. The tubes were spun down at 300 g for 5 minutes, then the supernatant was filtered through a 0.45 µm filter into 300 µL aliquots. Aliquots were stored at -80°C.

### Generating Stable Lines with Lentivirus

200, 000 cells were seeded per well in a 24 well plate. Polybrene was added (EMD-Millipore TR-1003-G) to a final concentration of 8 µg/ml in a total volume of 1mL in a well of a 24 well plate, then viral aliquots thawed at 37°C were then added dropwise into the well, then returned to the incubator. The next day, media was replaced. 48 hours after infection, antibiotic selection was started with 1 µg/mL puromycin until the control well died.

### Live Cell Confocal Microscopy

Adherent cells were plated (35,000 cells/well in an 8-well chamber slide) one day before small molecule treatment. Doxycycline-inducible lines were plated two days before small molecule treatment at half the seeding density. One day before treatment, cells are treated with doxycycline to a final working concentration of 1 μg/mL in 250 μL of complete growth media. On the day of treatment, wells are gently rinsed with 2 x 300 μL PBS, then incubated with 250 µl of complete phenol red-free growth media with small molecules added for the indicated time. After the indicated time, 1 µL of a fresh Hoechst 33342 solution (0.5 μg/μL in PBS) was added directly to the existing media in each well to achieve a final Hoechst 33342 concentration of 2 μg/mL, and incubated for 10 minutes. Cells were imaged with a Nikon A1R confocal microscope using a Plan Fluor 60X oil immersion, 1.30-numerical aperture objective. The microscope was equipped with a 405-nm violet laser, a 488-nm blue laser, a 561-nm green laser, and a 639-nm laser. Images were exported and analyzed using the FIJI software package.

### Fixed-Cell Confocal Microscopy

Adherent cells were plated (50,000 cells/well in an 8-well chamber slide) one day before the experiment. Cells were then washed three times with DPBS and fixed with 4% paraformaldehyde in PBS for 15 minutes at room temperature, washed three times, and permeabilized with 0.1% Triton for 5 minutes at room temperature. Cells were blocked in 10% goat serum in PBS for 1 hour at room temperature, and incubated with primary antibody overnight at 4°C. Cells were washed with PBS three times, then incubated with secondary antibody and DAPI for 1 hour at room temperature. Cells were washed with PBS and imaged with a Nikon A1R confocal microscope using a Plan Fluor 60X oil immersion 1.30-numerical aperture objective. The microscope was equipped with a 405-nm violet laser, a 488-nm blue laser, a 561-nm green laser, and a 639-nm laser. Images were exported and analyzed using the FIJI software package.

Custom Imaging Analysis Pipeline Construction:

1) **Segmentation**: A binary image is generated from a nuclear stained image (e.g. DAPI) of a field of view using Otsu thresholding. This binary image is then used as a mask to label coordinates within an acquired field of view corresponding to nuclei. Watershed transform is carried out to segment touching nuclei, and individual nuclei within a field of view are assigned with different numerical labels starting from 1 (“nlabels”).
2) **Generating masks**: “Celllabels” – “nlabels” are then expanded by 6 pixels circumferentially to capture cell area. At the same time, “nlabels” is also expanded by 3 pixels to generate “expandednlabels”. Cytoplasmic labels (“clabels”) are obtained by subtracting the “expandednlabels” from “celllabels”. “Nlabels” is also shrunk by 3 pixels to avoid taking in nuclear boundary measurements for the cytoplasm or nuclear readings. These pixel numbers can also be user defined.
3) **Gating**: Average nuclear stain intensities per area (“avg. nuc int”) are then tabulated for these individual nuclei, and a histogram is generated. Kernel density estimation is used to generate a probability density function (PDF) from this histogram. A normal distribution centered around the main peak of the PDF based on the full width at half maximum (FHWM) is then generated and overlayed. Assuming the population “avg nuc int” follows a normal distribution, the ends of the normal distributions are set as the gate bounds. Any nuclei without “avg nuc int” within the set gate bounds is gated out and labels for those nuclei and their respective cytoplasmic label are dropped to give “gatednlabels” and “gatedclabels”. Alternatively, if the user wishes to set fixed values for the gate bounds, this can also be passed in as an argument in the script.
4) **Summing intensities per cell**: Fluorophore images for this field of view are overlayed with the “gatednlabels” and “gatedclabels” to extract flurophore intensities tabulated for every single nuclear and cytoplasmic label. Average fluorophore intensites for each nuclei/cytoplasm label are also generated by dividing the intensity over the area of the label, to give a nuclear and cytoplasmic flurophore concentration (pixel intensity per area) per cell.

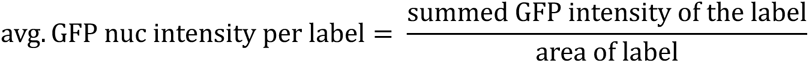
5) **Calculations and output.** For each cell, the average fluorophore pixel intensity per area is then used in the following calculation to give percent nuclear fluorophore:

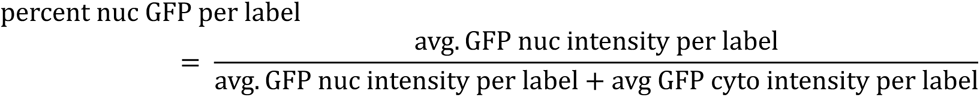 These values, including the average fluorophore intensity per label, are tabulated in a dataframe and output to an excel with a separate sheet for each condition. A summary of the mean for each value for each condition is also tabulated and output as an excel sheet. In addition, a folder is also created containing the gating histogram, as well as overlay masks onto the fluorophore images and the segmentation and nuclear masks from the original nuclear stain images for each field of view.

### Flow Cytometry

Adherent cells were lifted with trypsin and transferred into an Eppendorf or conical tube using complete media. Cells were pelleted at 300*g* and washed three times with PBS. Cells were resuspended in PBS and analyzed on either a BD Accuri C6+ or LSR II flow cytometer.

### Endogenous Tagging of Proteins Using Modified PITCh

To facilitate ease of cloning different target sgRNAs into the cassette, a modified all-in-one CRIPSR-Cas9 vector was made containing two orthogonal U6 promoter-stem loop pairs from the parental PITCh vector.^72^ The PITCH protocol was then followed to insert donor sequences into endogenous targets. Briefly, a day before transfection, adherent cells were seeded at a density of 25,000 cells in a 10 cm dish. On the day of transfection, media was replaced with 7 mL Opti-MEM. Both 1.2 µg of guide vector (containing target sgRNA, PITCh sgRNA and Cas9) and 0.6 µg of donor vector were added to 500 µl of Opti-MEM in an Eppendorf tube. To another Eppendorf tube containing 500 µl Opti-MEM was added 30 µl Lipofectamine 2000. Both Eppendorf tubes were gently mixed and left to incubate for 5 minutes at room temperature. The two solutions were then mixed and incubated for 30 minutes at room temperature, after which the mixture was added dropwise to the 10 cm plate containing cells. Cells were incubated for 24 hours, after which, the media was replaced with 10 mL of complete growth media. 72h after transfection, media was replaced with complete growth media containing puromycin (1 μg/ml), and selection continued by replacing media with fresh complete growth media supplemented with puromycin every two days. After complete death of control cells, puromycin selection was stopped and cells were expanded for immunoblotting.

### Immunoblotting for Endogenously Tagged Proteins

Cells were lifted with trypsin then washed with PBS three times and lysed with RIPA buffer supplemented with protease inhibitor cocktail (Roche) and 0.1% Benzonase (Millipore-Sigma) on ice for 15 minutes in Eppendorf tubes. The lysates were centrifuged at 21,000*g* for 15 minutes at 4 °C. The supernatant was collected, and the protein concentration was determined by BCA assay (Pierce). Equal amounts of lysates were loaded onto 4-12% Bis-Tris gel and separated by sodium dodecyl sulfate-polyacrylamide gel electrophoresis (SDS-PAGE). The gel was then transferred onto a nitrocellulose membrane and blocked with 5% milk in TBS-T for 1 hour at room temperature. The membrane was incubated with primary antibody overnight at 4 °C, and washed 3 times with TBS-T. Subsequently, the membrane was incubated with secondary antibody for 1 hour at room temperature and washed three times with TBS-T for visualization with an Odyssey CLx Imager (LI-COR).

### Sodium Arsenite Treatment and Imaging

Adherent cells were plated (25,000 cells/well in an 8-well chamber slide) two days before the experiment. One day before treatment, cells were treated with doxycycline to a final working concentration of 1 μg/mL in 250 μL of complete growth media. On the day of the experiment, cells were treated with 30 µM sodium arsenite in 250 μL complete phenol red-free media for an hour to induce stress granule formation. After an hour, small molecules solutions were added directly into the media in the well, and cells were incubated for another 3 hours. Cells were then washed three times with PBS and fixed with 4% paraformaldehyde in PBS for 15 minutes at room temperature, washed three times, and permeabilized with 0.1% Triton for 5 minutes on ice. Cells were blocked in 10% goat serum in PBS for 1 hour at room temperature then incubated with primary antibody overnight at 4°C. Cells were washed with PBS three times, then incubated with secondary antibody and DAPI for 1 hour at room temperature. Cells were washed with PBS and imaged with a Nikon A1R confocal microscope using a Plan Fluor 60X oil immersion 1.30-numerical aperture objective. The microscope was equipped with a 405-nm violet laser, a 488-nm blue laser, a 561-nm green laser, and a 639-nm laser. Images were exported and analyzed using the FIJI software package.

For continuous imaging, adherent cells were plated (25,000 cells/mL) in a 24 well glass bottom plate (CELLTREAT Scientific Products 229125) two days prior to treatment. One day before treatment, cells were treated with doxycycline to a final working concentration of 1 μg/mL in 500 μL of complete growth media.

On the day of imaging, media in the well was replaced with complete phenol red-free media containing 2 μg/mL Hoechst 33342 and 10 µM sodium arsenite. After an hour, the media in the wells was replaced with complete phenol red-free media containing 2 μg/mL Hoechst 33342, 10 µM sodium arsenite and the requisite concentration of small molecule. Immediately after treatment was administered, timelapse imaging was initiated. The plate was maintained at 37°C with 5% of CO_2_ in a stage top incubator (Okolab) over the course of 2.5 hours. Images were acquired by manually capturing images every 5 minutes at a fixed field of view. The Nikon A1R confocal microscope Plan Fluor 60X oil immersion 1.30-numerical aperture objective was used.

### Dorsal Root Ganglion (DRG) Neuron Harvesting and Culturing

All animals were housed and experiments were performed in accordance with the US National Institutes of Health (NIH) guidelines for the care and use of laboratory animals, and were approved by Stanford University’s Administrative Panel on Laboratory Animal Care. (APLAC # 20608). Two days before harvesting primary dorsal root ganglia cells (DRGs), 250 uL of poly-D lysine (PDL) was added to the bottom of an 8-well chamber slide or 24-well plate to coat the bottom of the well evenly. The slide was incubated at 37 °C in a sterile environment. The next day, PDL was removed, and the wells were washed with tissue culture-grade sterile water. The plate was left to dry with the lid open within a sterile biosafety cabinet for ∼ 1h. The slides were subsequently wrapped with parafilm then left in a 4 °C refrigerator.

#### Explant Harvesting

Primary dorsal root ganglia (DRG) cells were isolated from embryonic (day 17.5-18.5) Sprague-Dawley rats (Charles River Laboratories). A timed pregnant rat was euthanized with carbon dioxide and the embryos retrieved and kept in ice-cold Hank’s Buffered Salt Solution (HBSS, Corning 21-023-CM) supplemented with 10mM HEPES (Gibco 15630130). The embryo is decapitated, the ventral organs are removed and the body oriented with the dorsal side facing up. Under a dissection microscope, the skin and the spinal cord are removed and individual DRGs are extracted from either side of the cavity between spinal levels. The DRGs are kept in DMEM/10% FBS on ice until dissection of all embryos are complete (about 2-3h), aseptically transferred into a laminar flow hood, washed 5-10x with sterile HBSS-HEPES and digested with 0.25% trypsin (Gibco 15050065) supplemented with 80 Kunits of DNAseI (Sigma DN25) for 30 minutes. The trypsin is neutralized by adding an equal volume of DMEM/10% FBS and the cells are triturated via mechanical dissociation with a P200 pipette tip (∼15-30 times) and filtered through a 100μm cell strainer. Cells are spun down at 250*g* for 5 minutes and resuspended in complete medium, which consists of Neurobasal (Gibco 12348017) medium supplemented with 1% GlutaMAX (Gibco 35050061), 2% B-27 (Gibco 17504044), 1% penicillin/streptomycin, 2.5mg/mL D-glucose (Sigma G7021) and 50ng/mL NGF (2.5S beta subunit from Cedarlane Laboratories CLMCNET-001.25) that has been pre-equilibrated in a humidified 5% CO2 incubator at 37°C for 30 minutes. Single DRGs were seeded in the middle of a PDL coated well in the chamber slide in 15 μl of media. The slide or plate was left to incubate at 37 °C for 30 minutes in a 5% CO_2_ incubator to allow DRGs to adhere, then 250 µL of neurobasal supplemented with 40 µM of 5-fluoro-2’-deoxyuridine (FUDR, Sigma F0503) and NGF was carefully added to the well. Neurobasal media supplemented with FUDR and NGF was replaced every other day.

### AAV Production

On day 0, HEK293T cells were seeded in a 6-well plate for 70% confluency on Day 1. On Day 1, HEK293T cells were co-transfected with plasmids encoding the transgene, RepCap2, and adenovirus E4, E2A, and VA using the PEI MAX transfection reagent according to the manufacturer’s protocol. The morning of Day 2, media was replaced with complete DMEM. On Day 3, the viral supernatant was removed and filtered through a 0.45 µm filter.

### DRG Neuron Infection with AAV

Two days post explant seeding, media in the wells were replaced with 250 μL 0.2 µm filtered neurobasal media supplemented with FUDR (40μM) and NGF (50 ng/mL). 8 μL of NMNAT1 transgene viral supernatant and 12 μL of eDHFR transgene viral supernatant was added to each well. 4 days post-infection, neurons were examined for transgene expression.

### Axotomy and Neuron Imaging

Prior to the cut, fluorescence images of entire explants were acquired in an epi-fluorescence microscope (Leica DMI 6000B) using a 10X (0.32 NA) objective. The microscope is equipped with ORCA-Flash4.0 Digital CMOS camera, Lumencor SOLA light source, and filter sets: 370-39/409/448-63 nm (blue emission), 484-25/505/524-32 nm (green emission), 560-32/581/607-40 nm (red emission), and 640-19/655/680-30 nm (far-red emission).

Biopsy punch (1.0 mm, Royaltek) was sterilized with 70% EtOH, then dipped into neurobasal media. The biopsy punch was used to severe the cell body from the neurites. The detached cell body was then extracted with sterilized tweezers. To ensure complete removal of the cell body, fluorescence images of entire explants were acquired again in an epi-fluorescence microscope (Leica DMI 6000B) using 10x (0.32 NA) objective post-cut.

To observe degeneration of the neurites, axon termini and the proximal axon region were imaged with a Nikon A1R confocal microscope using a Plan Fluor 60X oil immersion at 0h, 3h, 6h, 12h, 24h, 48h, 72h, 96h, post injury.

### Synthetic Chemistry

All synthetic chemistry procedures are available in the Supplementary Information.

### DNA Sequences

All DNA sequences used can be found in the Supplementary Information.

