## Extended Data for "Targeted Protein Relocalization via Protein Transport Coupling"

### Extended Data Figures and Figure Captions:

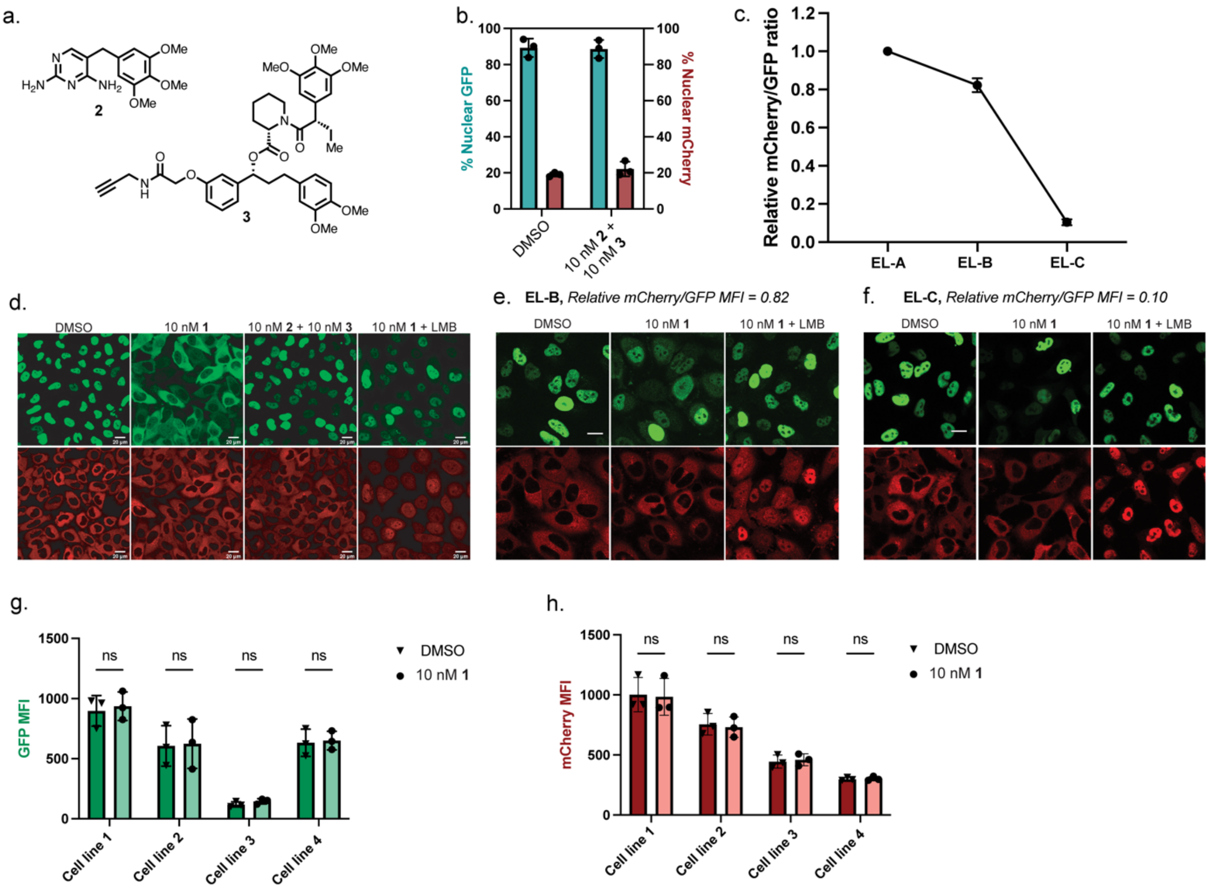


**Extended Data Fig. 1: Examination of NMNAT1 export cell lines. a**, Unlinked warhead controls for **1. 2:** trimethoprim (TMP), binds ecDHFR, **3:** binds FKBP12$^{F36V}$. **b**, No change of localization of either protein upon treatment of the export line with unlinked warheads **2** and **3**. **c**, Comparison of the relative mCherry/GFP median fluorescence intensity ratio between the three isolated clonal export lines (**EL-A-C**). **d**, Representative live-cell images for NMNAT1 relocalization treated with **1** or control molecules, or **1** and leptomycin-B (LMB) in Export Line A (EL-A) after 3 hour treatment. **e**, Representative live-cell images for NMNAT1 relocalization treated with **1** or control molecules, or **1** and leptomycin-B (LMB) in Export Line B (**EL-B**) after 3 hour treatment. **f**, Representative live-cell images for NMNAT1 relocalization treated with **1** or control molecules, or **1** and leptomycin-B (LMB) in Export Line C (EL-C) after 3 hour treatment. **g**, Levels of GFP after treatment with **1** for 3 hours across four isolated export cell lines with varying nuclear protein expression levels. **h**, Levels of mCherry after treatment with **1** for 3 hours across four isolated export cell lines. MFI : Mean fluorescence intensity. Images in **d**, **e**, **f** are representative of three biological replicates. Data are compiled from three independent experiments. Scale bars are 20 μm.

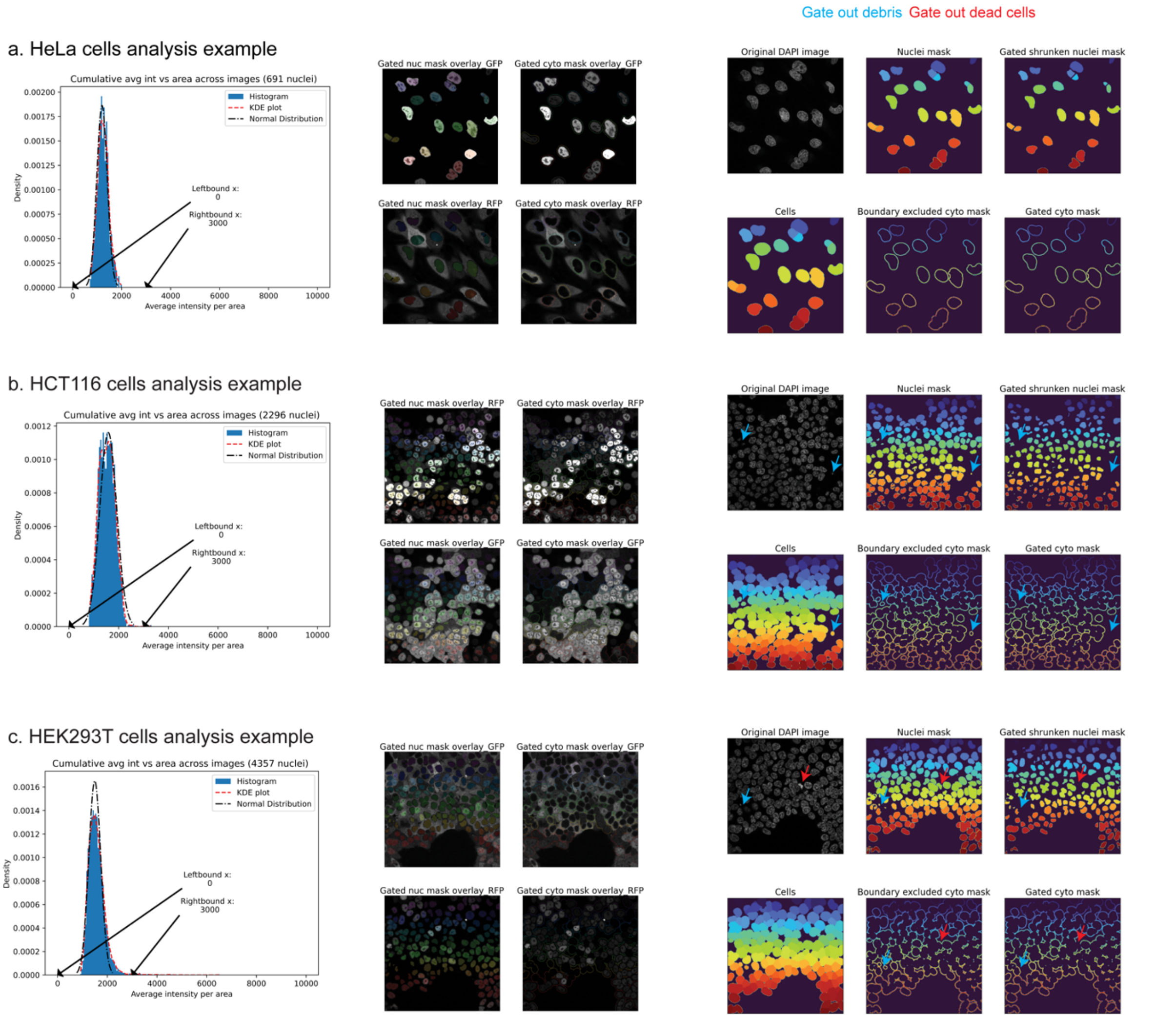


**Extended Data Fig. 2: Illustration of gating and segmentation pictures automatically generated from analysis pipeline applied to three different cell types. a**, Computational pipeline applied on HeLa cells. Gating histograms generated with the set bounds indicated. Masks for nuclear and cytoplasm areas overlayed onto respective fluorophore images. Cytoplasm areas are circumferentially grown from nuclei masks retaining the relative size between the masks. **b**, Computational pipeline applied on HCT116 cells. Masks for nuclear and cytoplasm areas overlayed onto respective fluorophore images with segmentation of highly confluent cells using thin cytoplasm masks. The nuclear and cytoplasm masks corresponding to debris (blue arrows) are gated out. **c**, Computational pipeline applied on HEK293T cells. Masks for nuclear and cytoplasm areas overlayed onto respective fluorophore images, with segmentation of highly confluent cells and gating out of dead cells (high nuclear stain over a small area) and debris.

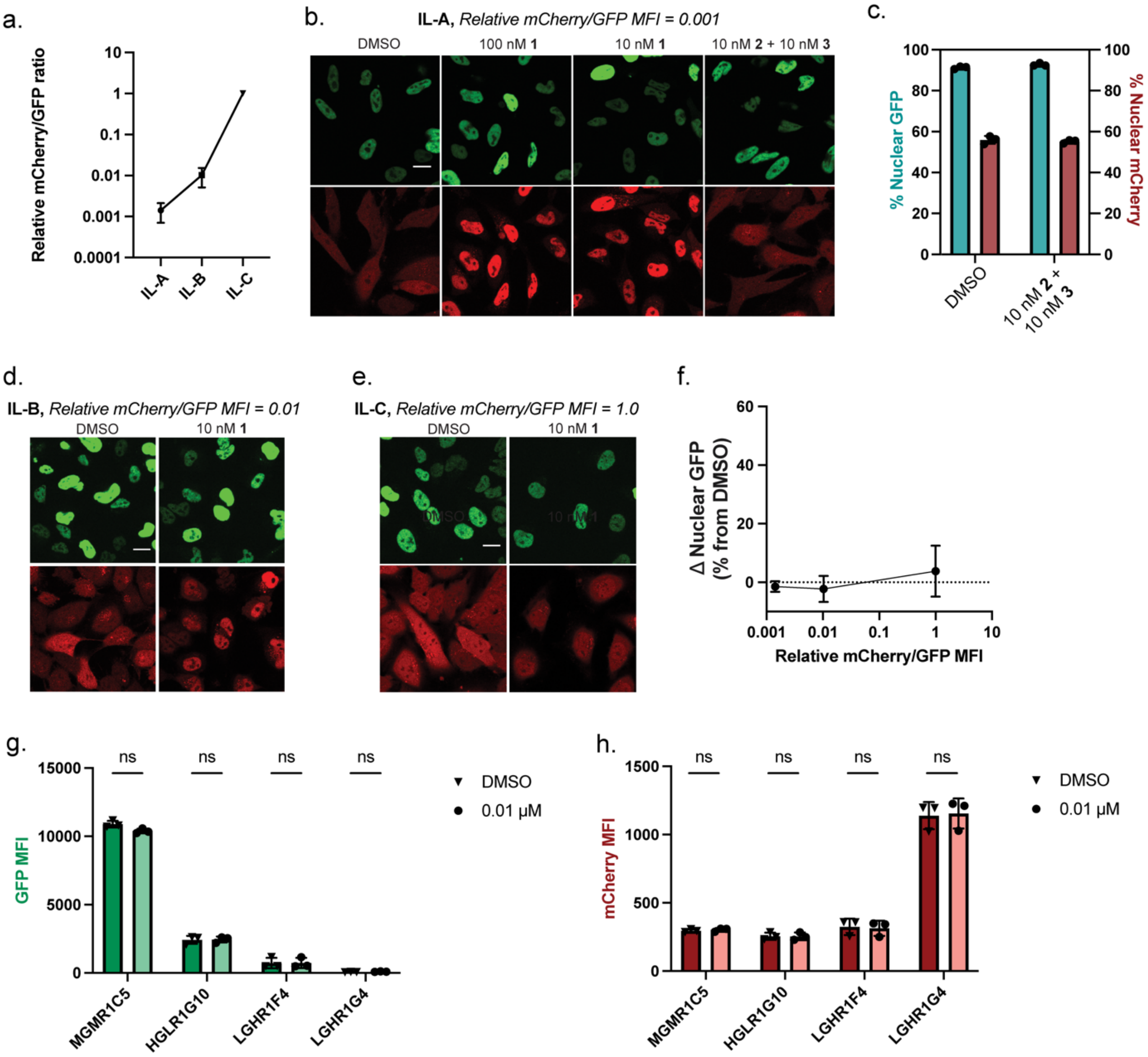


**Extended Data Fig. 3: Examination of NMNAT1-utilizing import cell lines.** **a**, Comparison of the relative mCherry/GFP median fluorescence intensity ratio between the three isolated clonal import lines (**IL-A-C**). **b**, Representative live-cell images of clonal import line A (IL-A) upon treatment with **1** or small molecule controls for 3 hours. **c**, Quantification of localization upon treatment of the IL-A with unlinked warheads **2** and **3. d**, Representative live-cell images of clonal import line B (IL-B) upon treatment with **1** for 3 hours. **e**, Representative live-cell images of clonal import line C (IL-C) upon treatment with **1** for 3 hours. **f**, Quantification of NMNAT1 protein localization upon treatment with 10 nM **1** in IL-A-C possessing different mCherry/GFP ratios. **g**, Levels of GFP after treatment with **1** for 3 hours in four isolated import cell lines with varying nuclear protein expression levels. **h**, Levels of mCherry after treatment with **1** for 3 hours across four isolated import cell lines. Images in **b**, **d, e,** are representative of three biological replicates. Data is compiled from three independent experiments. Scale bars are 20 μm. MFI = median fluorescence intensity.

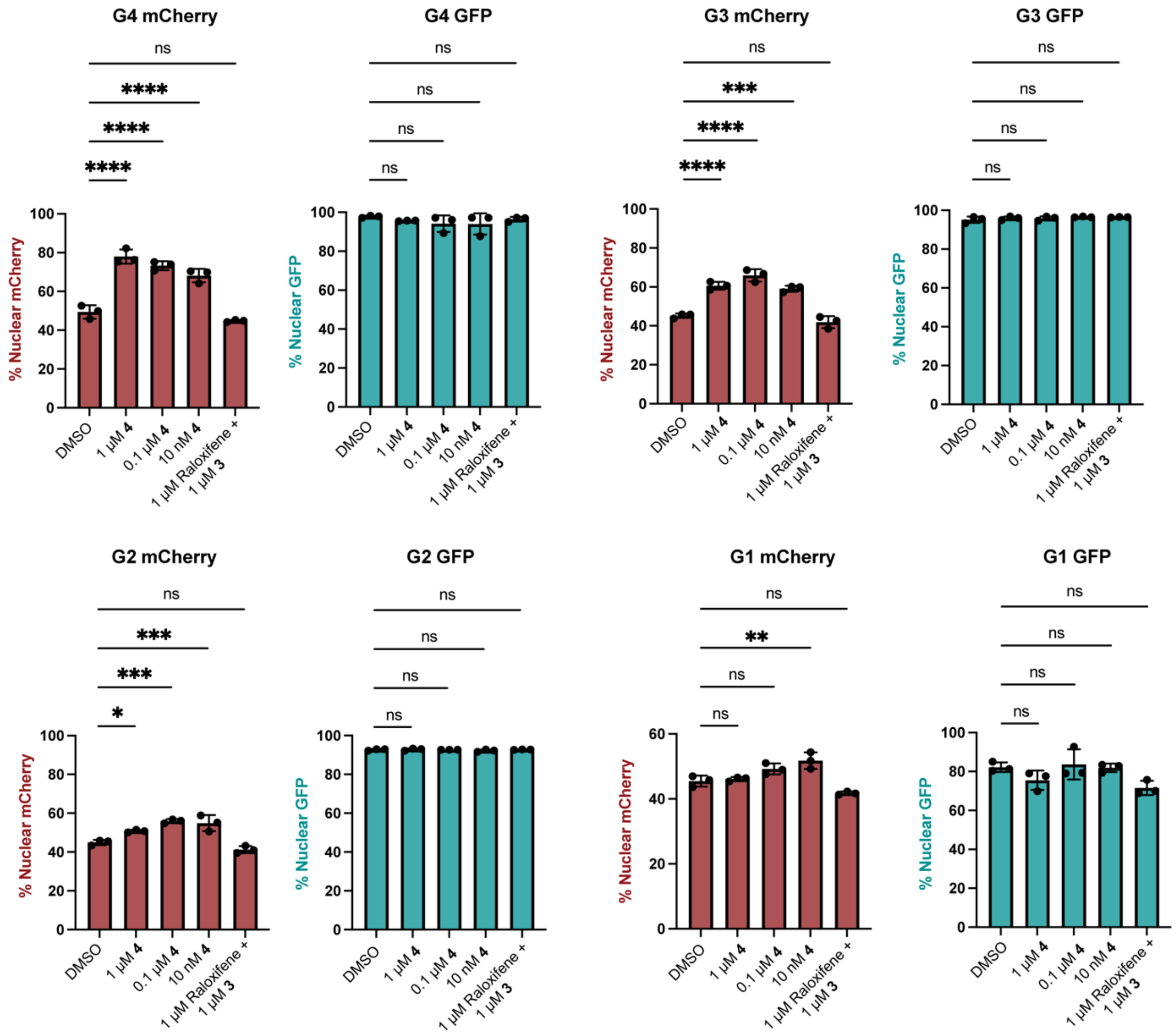


**Extended Data Fig. 4: Comparisons of percentage nuclear fluorophores across different treatment conditions in cells grouped based on ERα-GFP expression for SMARCB1$^{Q318X}$ HeLa cells.**

Data is compiled from three independent experiments. P values were determined by one-way ANOVA comparing each condition to the DMSO control. P values: (*) indicates P ≤ 0.05 (**) indicates P ≤ 0.01, (***) indicates P ≤ 0.001, P ≤ 0.0001. Cells were divided into four groups based on their average nuclear GFP intensity G4: 4500 > x ≥ 4500, G3: 3000 > x ≥ 1500, G2: 1500 > x ≥ 500, G1: 500 > x ≥ 50.

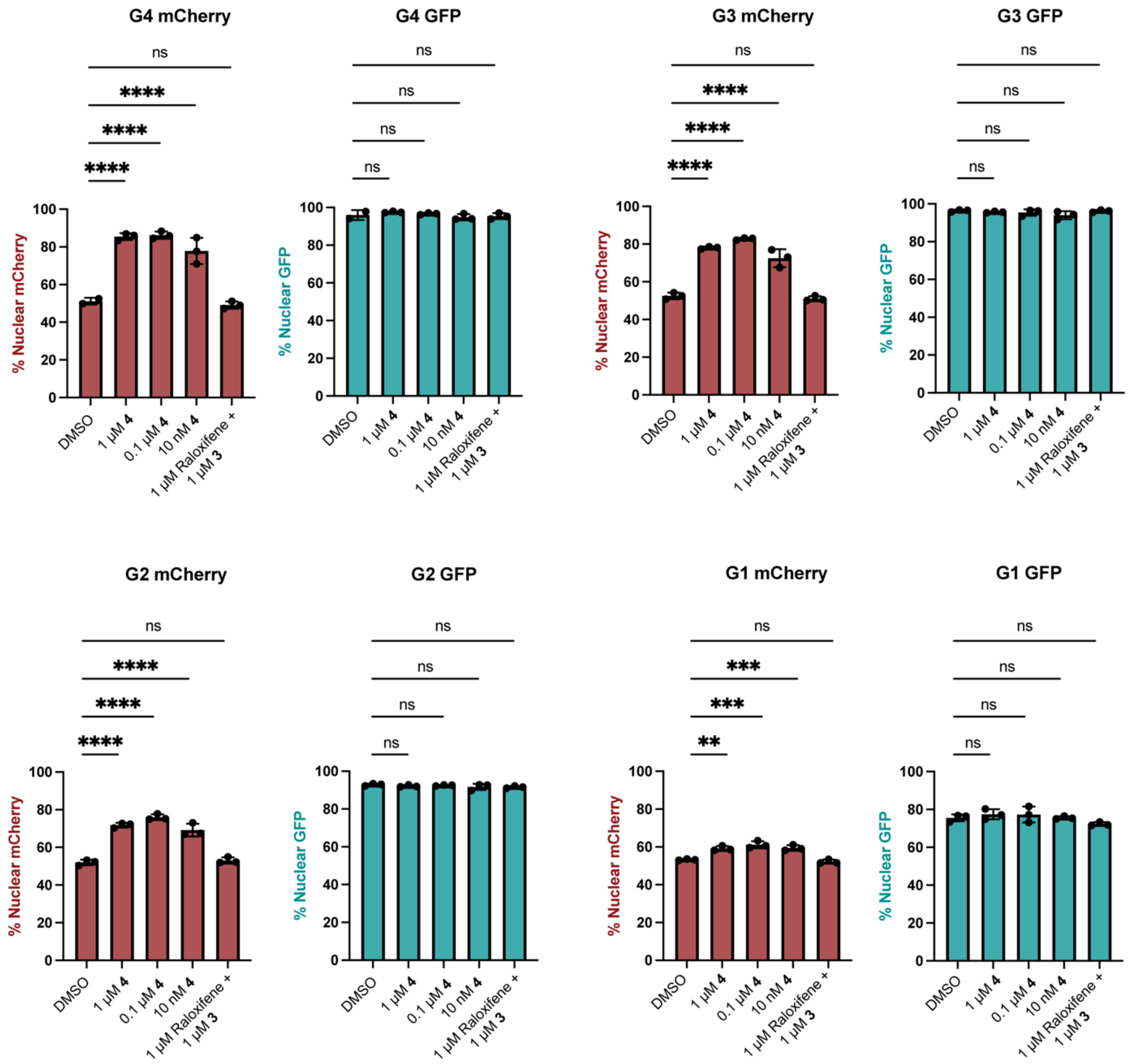


**Extended Data Fig. 5: Comparisons of percentage nuclear fluorophores across different treatment conditions in cells grouped based on ERα-GFP expression for TDP43$^{\Delta NLS}$ HeLa cells.**

Data is compiled from three independent experiments. P values were determined by one-way ANOVA comparing each condition to the DMSO control. P values: (*) indicates P ≤ 0.05 (**) indicates P ≤ 0.01, (***) indicates P ≤ 0.001, P ≤ 0.0001. Cells were divided into four groups based on their average nuclear GFP intensity G4: 4500 > x ≥ 4500, G3: 3000 > x ≥ 1500, G2: 1500 > x ≥ 500, G1: 500 > x ≥ 50.

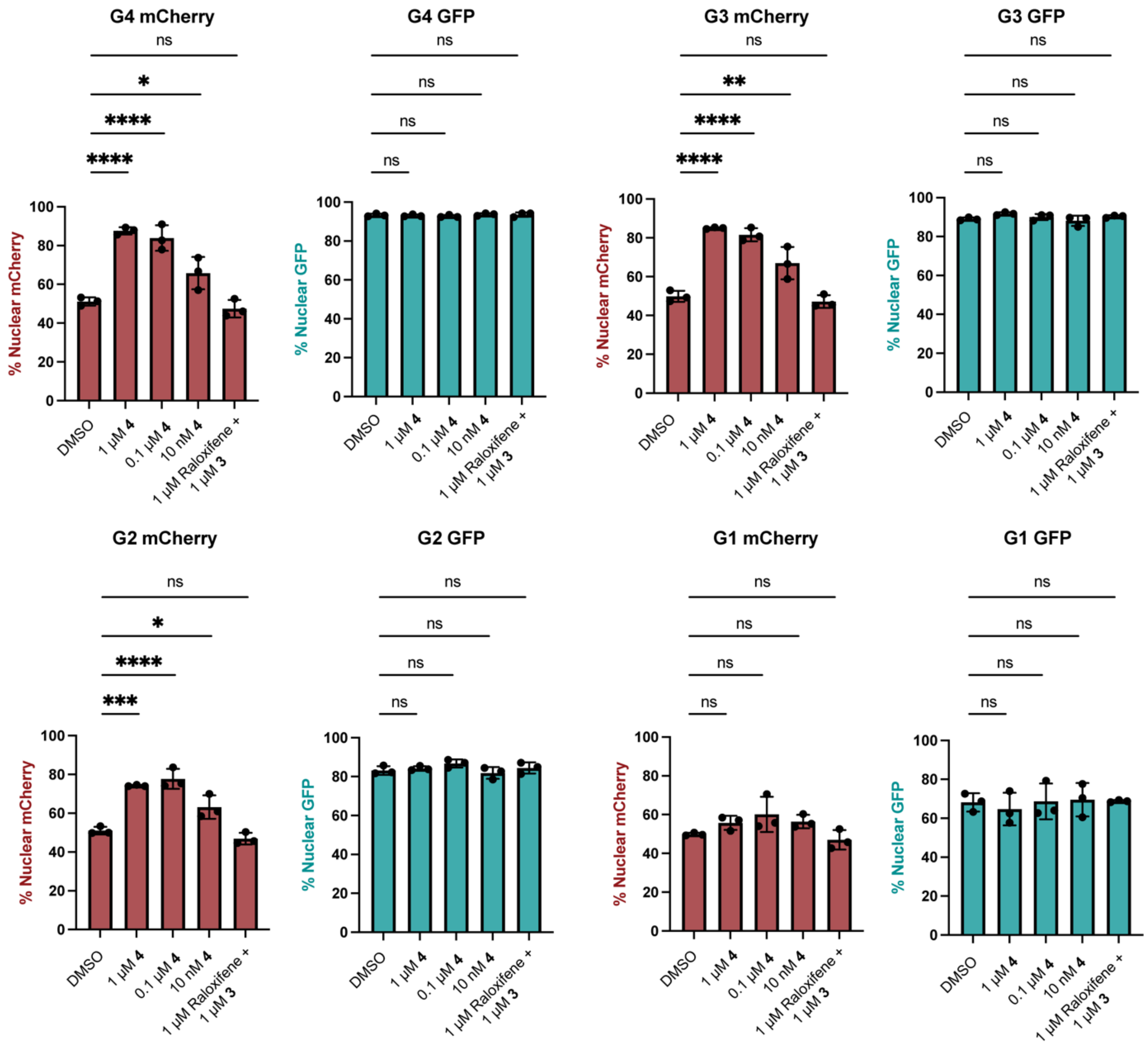


**Extended Data Fig. 6: Comparisons of percentage nuclear fluorophores across different treatment conditions in cells grouped based on ERα-GFP expression for $FUS^{R495X}$ HeLa cells.**

Data is compiled from three independent experiments. P values were determined by one-way ANOVA comparing each condition to the DMSO control. P values: (*) indicates $P \leq 0.05$ (**) indicates $P \leq 0.01$, (***) indicates $P \leq 0.001$, $P \leq 0.0001$. Cells were divided into four groups based on their average nuclear GFP intensity G4: $1500 > x \geq 800$, G3: $800 > x \geq 500$, G2: $500 > x \geq 200$, G1: $200 > x \geq 50$.

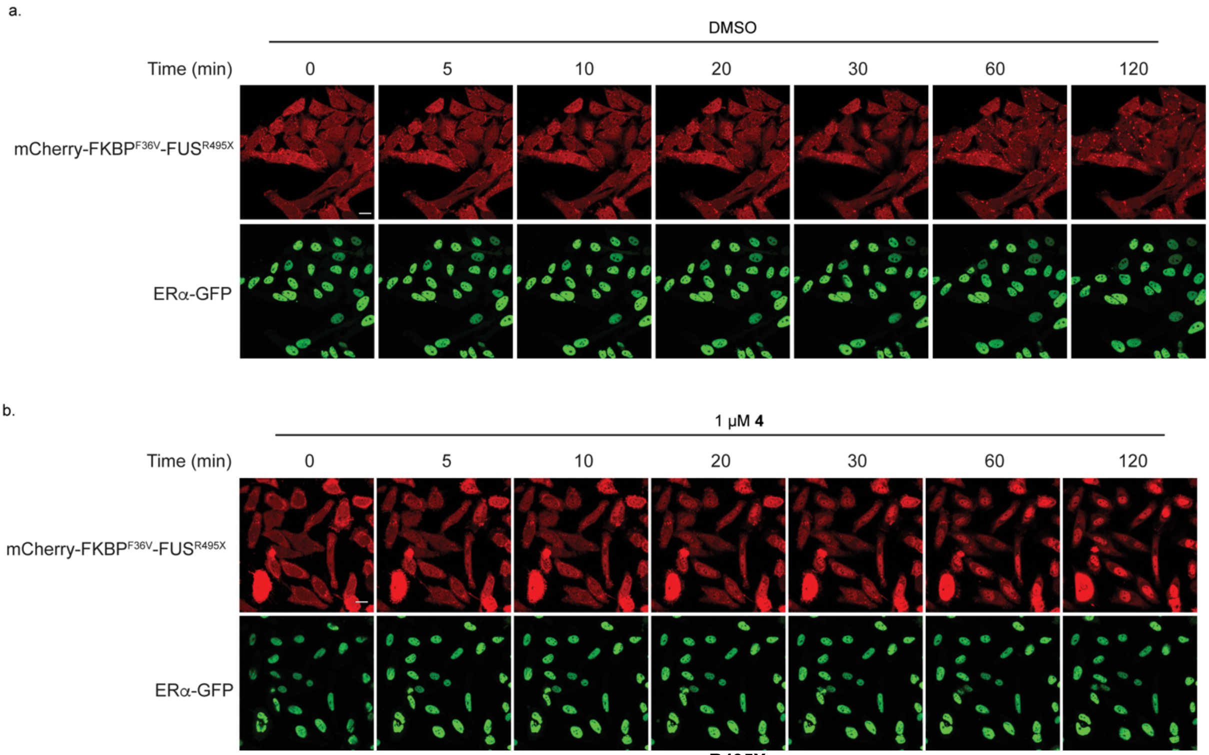


**Extended Data Fig. 7: Timelapse snapshots of FUS$^{R495X}$ extraction from granules.**
**a**, Timelapse imaging snapshots of FUS$^{R495X}$ extraction from granules when cells are treated with DMSO**. b**, Timelapse imaging snapshots of FUS$^{R495X}$ extraction from granules when cells are treated with **4**. Images are representative of two independent experiments. Scale bars are 20 μM.

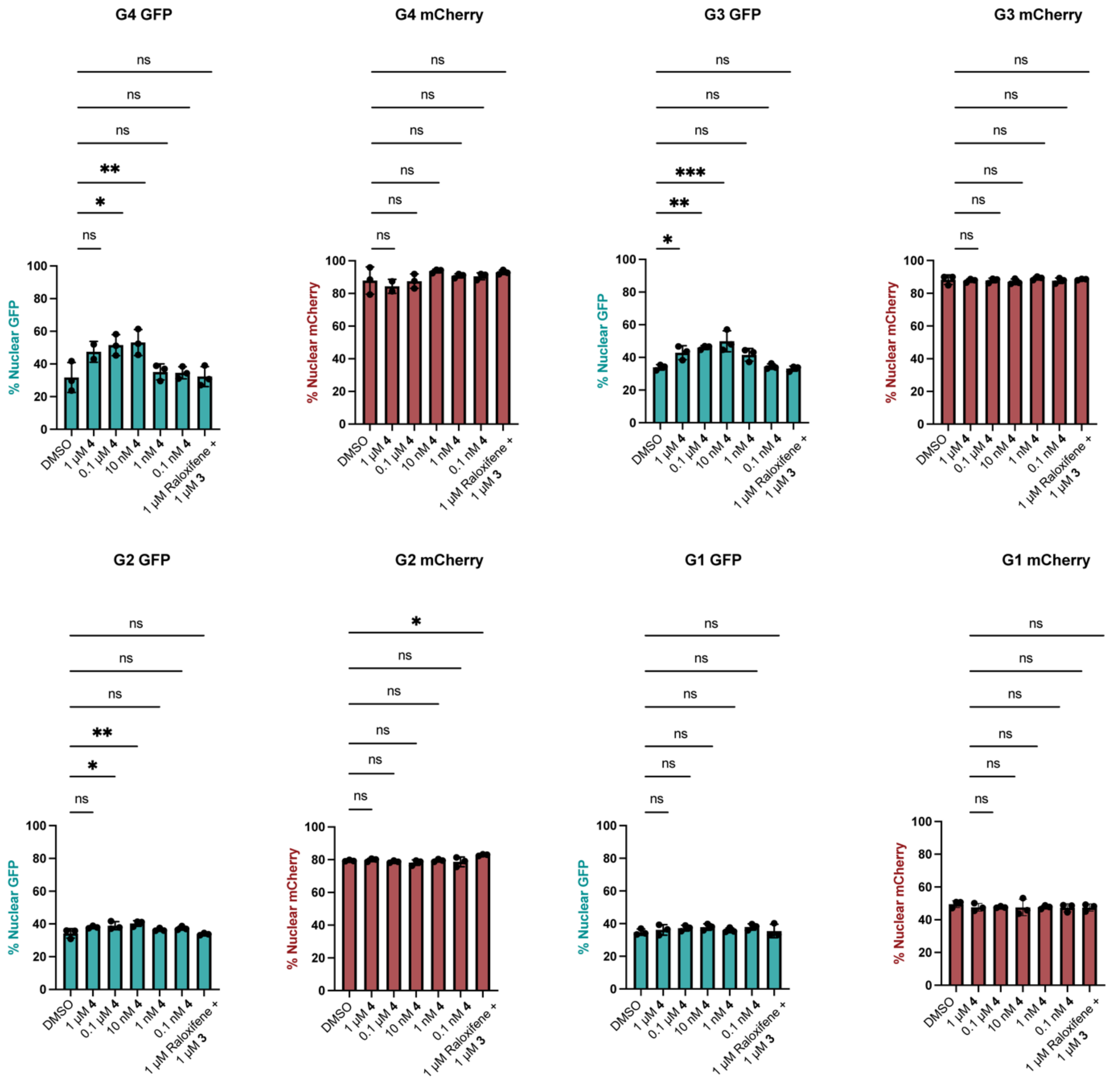


**Extended Data Fig. 8: Comparisons of percentage nuclear fluorophores across different treatment conditions in cells grouped based on ERα-mCherry expression for HEK293T FOXO3A-GFP-FKBP12$^{F36V}$ knock-in cells.**

Data is compiled from three independent experiments. P values were determined by one-way ANOVA comparing each condition to the DMSO control. P values: (*) indicates $P \leq 0.05$ (**) indicates $P \leq 0.01$, (***) indicates $P \leq 0.001$, $P \leq 0.0001$. Cells were divided into four groups based on their average nuclear mCherry intensity G4: $4900 > x \geq 2000$, G3: $2000 > x \geq 800$, G2: $800 > x \geq 200$, G1: $200 > x \geq 50$.

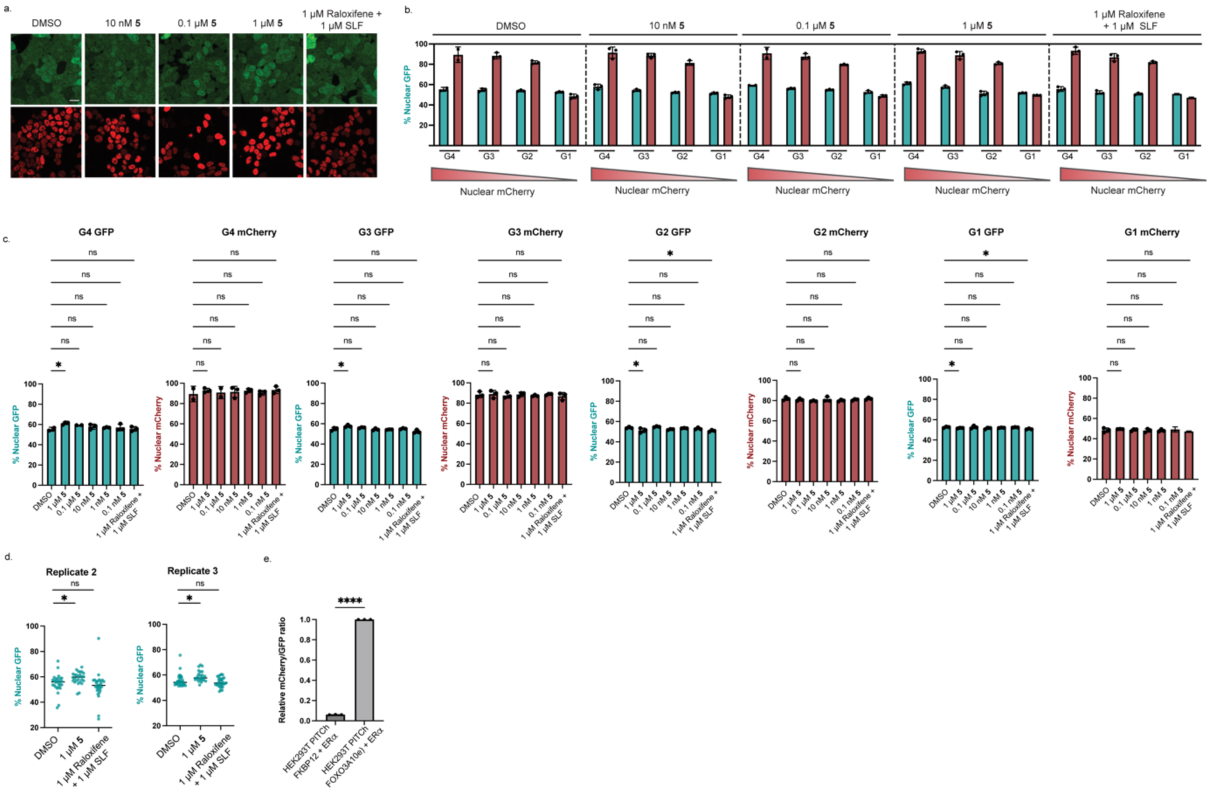


**Extended Data Fig. 9: Import of endogenous FKBP12 by ERα in HEK293T cells**

**a**, Representative live-cell images of HEK293T cells stably expressing ERα-mCherry after treatment with **5** or small molecule controls for 3 hours. **b**, Group quantitative analysis of FKBP12 knock-in HEK293T treated with varying concentrations of **5** for 3 hours. Cells were divided into four groups based on their average nuclear mCherry intensity G4: $4900 > x \geq 1800$, G3: $1800 > x \geq 800$, G2: $800 > x \geq 200$, G1: $200 > x \geq 50$. **c**, Comparisons of percentage nuclear fluorophores across different treatment conditions, with cell groupings as in b. P values were determined by one-way ANOVA comparing each condition to the DMSO control. **d**, Quantitative analysis of nuclear FKBP12 in the top 5% ER-expressing HEK293T cells, biological replicate 2, number of cells: DMSO: 27, 1 µM **7**: 26, 1 µM Raloxifene + 1 µM SLF: 27. Biological replicate 3, number of cells: DMSO: 29, 1 µM **7**: 24, 1 µM Raloxifene + 1 µM SLF: 30. **e**, Relative mCherry/GFP median fluorescence intensity comparison between the FKBP12 knock-in HEK293T line and the FOXO3A knock-in HEK293T line. Scale bars are 20 µm. P values: (*) indicates $P \leq 0.05$ (**) indicates $P \leq 0.01$, (***) indicates $P \leq 0.001$, $P \leq 0.0001$. Images in **a** are representative of three biological replicates. Data in **b**, **c** are compiled from three independent experiments.

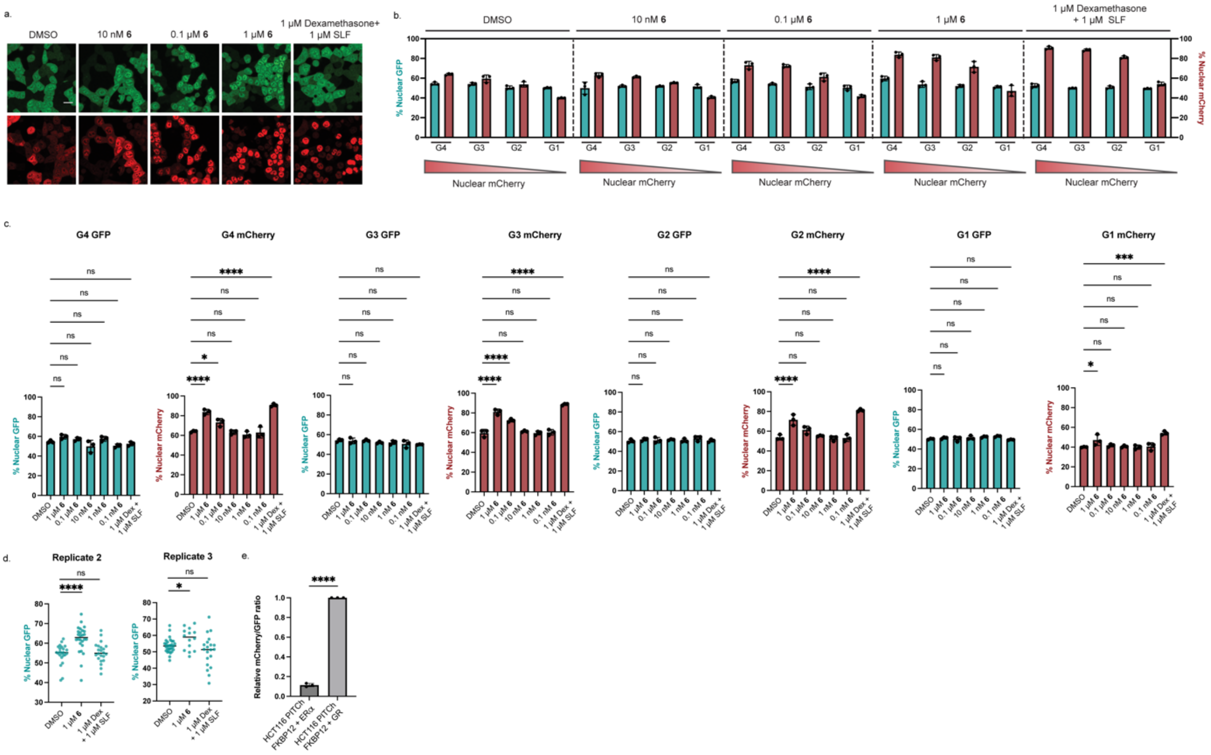


**Extended Data Fig. 10: Import of endogenous FKBP12 by GR in HCT116 cells.**
**a**, Representative live-cell images of HCT116 cells stably expressing GR-mCherry after treatment with **6** or small molecule controls for 3 hours. **b**, Group quantitative analysis of FKBP12 knock-in HCT116 cells treated with varying concentrations of **6** for 3 hours. Cells were divided into four groups based on their average nuclear mCherry intensity G4: 4900 > x ≥ 2000, G3: 2000 > x ≥ 800, G2: 800 > x ≥ 200, G1: 200 > x ≥ 50. **c**, Comparisons of percentage nuclear fluorophores across different treatment conditions, with cell groupings as in b. P values were determined by one-way ANOVA comparing each condition to the DMSO control. **d**, Quantitative analysis of nuclear FKBP12 in the top 5% GR-expressing HCT116 cells, biological replicate 2, number of cells: DMSO: 23, 1 μM **7**: 26, 1 μM Raloxifene + 1 μM SLF: 22. Biological replicate 3, number of cells: DMSO: 29, 1 μM **7**: 14, 1 μM Raloxifene + 1 μM SLF: 20. **e**, Relative mCherry/GFP median fluorescence intensity comparison between the FKBP12 knock-in HCT116 with ERα and FKBP12 knock-in HCT116 with GR. Scale bars are 20 μm. P values: (*) indicates P ≤ 0.05 (**) indicates P ≤ 0.01, (***) indicates P ≤ 0.001, P ≤ 0.0001. Images in **a** are representative of three biological replicates. Data in **b**, **c** are compiled from three independent experiments.

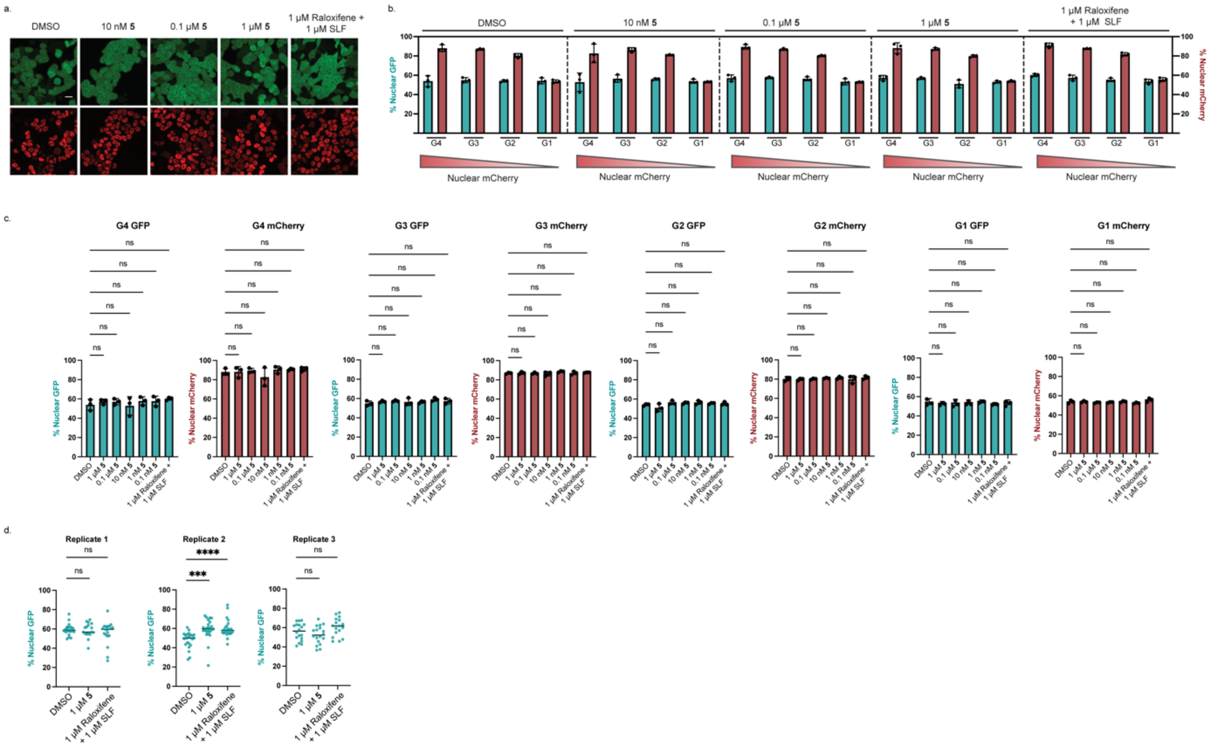


**Extended Data Fig. 11: Import of endogenous FKBP12 by ERα in HCT116.**

**a**, Representative live-images of HCT116 cells stably expressing ERα-mCherry **5** or small molecule controls for 3 hours. **b**, Group quantitative analysis of FKBP12 knock-in HCT116 treated with varying concentrations of **5** for 3 hours. Cells were divided into four groups based on their average nuclear mCherry intensity G4: $2500 > x \geq 800$, G3: $800 > x \geq 500$, G2: $500 > x \geq 200$, G1: $200 > x \geq 50$. **c**, Comparisons of percentage nuclear fluorophores across different treatment conditions, with cell groupings as in b. **d**, Quantitative analysis of nuclear FKBP12 in the top 5% ERα-expressing HCT116 cells. Scale bars are 20 µm. P values: (*) indicates P ≤ 0.05 (**) indicates P ≤ 0.01, (***) indicates P ≤ 0.001, P ≤ 0.0001. Images in **a** are representative of three biological replicates. Data in **b**, **c** are compiled from three independent experiments.

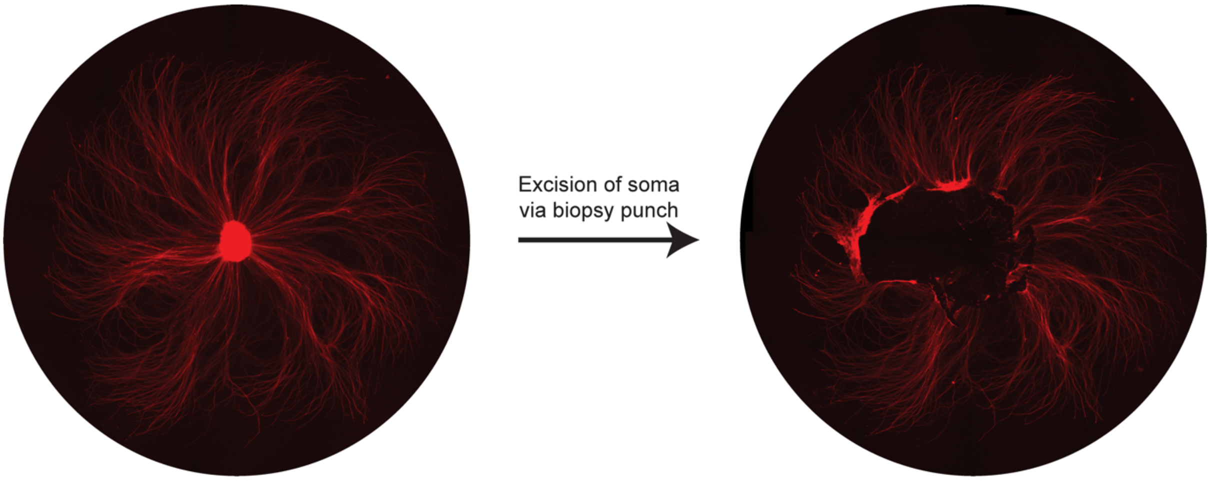


**Extended Data Fig. 12**: **Representative images of before and after axotomy on explants.** Axotomies were performed using a biopsy punch, the mRuby3-Axontag protein serves as an axonal marker.
